# Single Cell Proteomics Using a Trapped Ion Mobility Time-of-Flight Mass Spectrometer Provides Insight into the Post-translational Modification Landscape of Individual Human Cells

**DOI:** 10.1101/2022.02.12.480144

**Authors:** Benjamin C. Orsburn, Yuting Yuan, Namandjé N. Bumpus

## Abstract

Single cell proteomics is a powerful tool with potential for markedly enhancing understanding of cellular processes. Previously reported single cell proteomics innovations employ Orbitrap mass spectrometers. In this study we describe the development, optimization, and application of multiplexed single cell proteomics to the analysis of human-derived cells using trapped ion mobility time-of-flight mass spectrometry. This method, denoted as pasefRiQ is an advance as it allows accurate peptide quantification at picogram peptide concentrations. When employing a peptide carrier channel to boost protein sequence coverage, we obtain over 40,000 tandem mass spectra in 30 minutes, while achieving higher sequence coverage of each identified protein than described for SCOPE2. Using NCI-H-358 cells, which are a human bronchioalveolar carcinoma and KRAS^G12C^ model cell line, we demonstrate that the level of coverage achieved using this method enables the quantification of up to 1,255 proteins per cell and the detection of multiple classes of post-translational modifications in single cells. Further, when cells were treated with sotorasib, a KRAS^G12C^ covalent inhibitor, pasefRiQ revealed cell-to-cell variability in the impact of the drug on the NCI-H-358 cells, providing insight missed by traditional analyses. We provide multiple resources necessary for the application of single cell proteomics to drug treatment studies including tools to reduce cell cycle linked proteomic effects from masking pharmacological phenotypes.

**Significance Statement:** This work describes the establishment of a single cell proteomics method using a time-of-flight mass spectrometer. Through this approach, we demonstrate the confident identification of post-translational modifications in single human-derived cells. Additionally, using a KRAS^G12C^ covalent inhibitor as a model compound we show that this method can be used to understand pharmacological responses of single human-derived cultured cells.

## Introduction

Single cell RNA sequencing (scSeq) has markedly advanced understanding of biology at the level of individual cells.^1^ While an unquestionably powerful tool, a major limitation of all RNA based technology is the lack of correlation between transcript abundance and protein abundance in human systems.^2^ In addition, cellular function is often influenced by protein processing events such as proteolytic cleavage or post translational modifications (PTMs).^3^ As such, the direct analysis of the protein themselves in single cells is a promising avenue of research.^4^

Single cell proteomics using liquid chromatography mass spectrometry (LCMS) is a newly emerging field led by parallel improvements in both instrumentation and methodology.^5^ The majority of studies published to date have focused on the essential method development and proof of concept work necessary to set the stage for applications of the technology. Some of the most promising biological works described to date have utilized multiplexing reagents to obtain quantitative proteomic data on multiple cells in each single LCMS experiment. Multiplexing has several advantages, most notably allowing enough cells to be analyzed per study for sufficient statistical power to draw biological conclusions. In recent works, independent teams using these approaches have demonstrated the ability to study biological systems with multiplexed single cell proteomics including macrophage differentiation and diversity in cancer cell line populations.^6, 7^

Today single cell proteomics has demonstrated the ability to quantify hundreds of proteins per cell, largely driven by quantifying a relatively small number of peptides per protein. While accurate quantification of proteins can be derived from measurements of individual peptides, higher sequence coverage is required for the identification of many protein features. For example, PTMs such as phosphorylation and acetylation are only detected in proteomics studies where high relative sequence coverage is obtained or offline chemical enrichment is performed.^8, 9^

To date, all multiplex single cell proteomics studies have utilized various iterations of hybrid Orbitrap mass spectrometers (MS).^10, 11^ Orbitraps are popular MS systems due to their relatively high mass accuracy and resolution, characteristics that are largely obtained at the consequence of relative scan acquisition rate compared to other mass analyzers.^12^ In contrast, Time of Flight (TOF) systems are characterized by higher relative scan acquisition rates, and correspondingly lower resolution and mass accuracy. New advances in ion accumulation prior to TOF have successfully circumvented these traditional limitations by allowing new equilibriums between sensitivity and speed to be leveraged in LCMS workflows.^13^ In this study we explore the capabilities of a third generation trapped ion mobility time of flight mass spectrometer (TIMSTOF Flex) for sensitive and accurate multiplexed proteomics through a combination of parallel accumulation serial fragmentation for reporter ion quantification (pasefRiQ). As an application of pasefRiQ, we describe the analysis of single cells from the KRAS^G12C^ model lung cancer cell line NCI-H-358 (H358). We find that the number of proteins quantified in individual cells are sufficient to cluster each cell by their relative cell cycle stage. Importantly, we describe the first observations of multiple classes of protein PTMs by LCMS in single human cells and find that the quantification values for PTMs of well-studied proteins correlate with other phenotypic markers.

To explore the application of single cell proteomics in the context of molecular pharmacology, we treated cells with the KRAS^G12C^ covalent inhibitor, sotorasib.^14^ We find that when the proteomes of individual cells are handled as biological replicates during data analysis, sotorasib treatment largely mimics the effects observed in studies based on the proteomics of bulk cell homogenates. Single cell proteomics provides additional insight into these systems by allowing us to directly elucidate the cell-to-cell heterogeneity in response to inhibitor treatment. With this additional data we find evidence that some of the proteins displaying the largest differential response to sotorasib are disproportionately impacted in a relatively small number of individual cells. Taken together, these results demonstrate a powerful role for single cell proteomics in understanding cell-to-cell variability including in drug response

## Results and Discussion

### Practical intrascan linear dynamic of pasefRiQ across three orders of dynamic range

Due to the time of flight effect of fragment ions leaving the collision cell of the TIMSTOF analyzer, two MS2 scan events must be combined to obtain fragments from both high and low relative mass-to-charge ratios. By separately optimizing the pre-pulse storage time and collision energies of two trapping events pasefRiQ can provide optimal fragmentation for both peptide sequencing and maximum reporter ion signal for quantification (**S. Fig. 1**.)

A major historical challenge in protein mass spectrometry is the wide intracellular distribution of protein dynamic range which has been estimated to approximately 7-orders mammalian cells^15, 16^, which is a stark contrast to mass analyzers which may only have a two-order intrascan linear dynamic range^17^Limitations in dynamic range effect both our ability to detect lower abundance proteins of interest and to accurately quantify proteins exhibiting high relative fold change alterations between conditions.^18^ To evaluate the practical intrascan linear dynamic range of pasefRiQ we prepared a 4-order dilution series of a commercial K562 cell line digest with TMTPro 9-plex reagent. The mean reporter signal for each channel from all identified peptides maintained near linearity across the entire dilution series, with an R^2^ of 0.982 (**Fig. 1A**). While the number of missing reporter ions increases as the peptide concentration decreases, we do not observe a marked increase in the number of missing reporter values until we exceed an intrascan dilution of three orders. In the reporter ion channel that contained 200 picograms of K562 digest standard compared to a channel of 200 nanograms, the number of reporter ions detected decreased by 53.2%. Surprisingly, not all reporter ion signal was lost at even 10-fold below this level as 10.2% of peptides contained reporter ion signal at a level approximating 10 picograms on column (**Fig. 1B, S. Fig 2D**). These data suggested that pasefRiQ has an effective intrascan linear dynamic range of approximately 3-orders, which is substantially higher than Orbitrap instruments^17^. These results indicate pasefRiQ may not demonstrate the same limitations and subsequent ratio distortions recently described as the “carrier proteome effect” as Orbitrap systems.^19, 20^

**Fig. 1.**
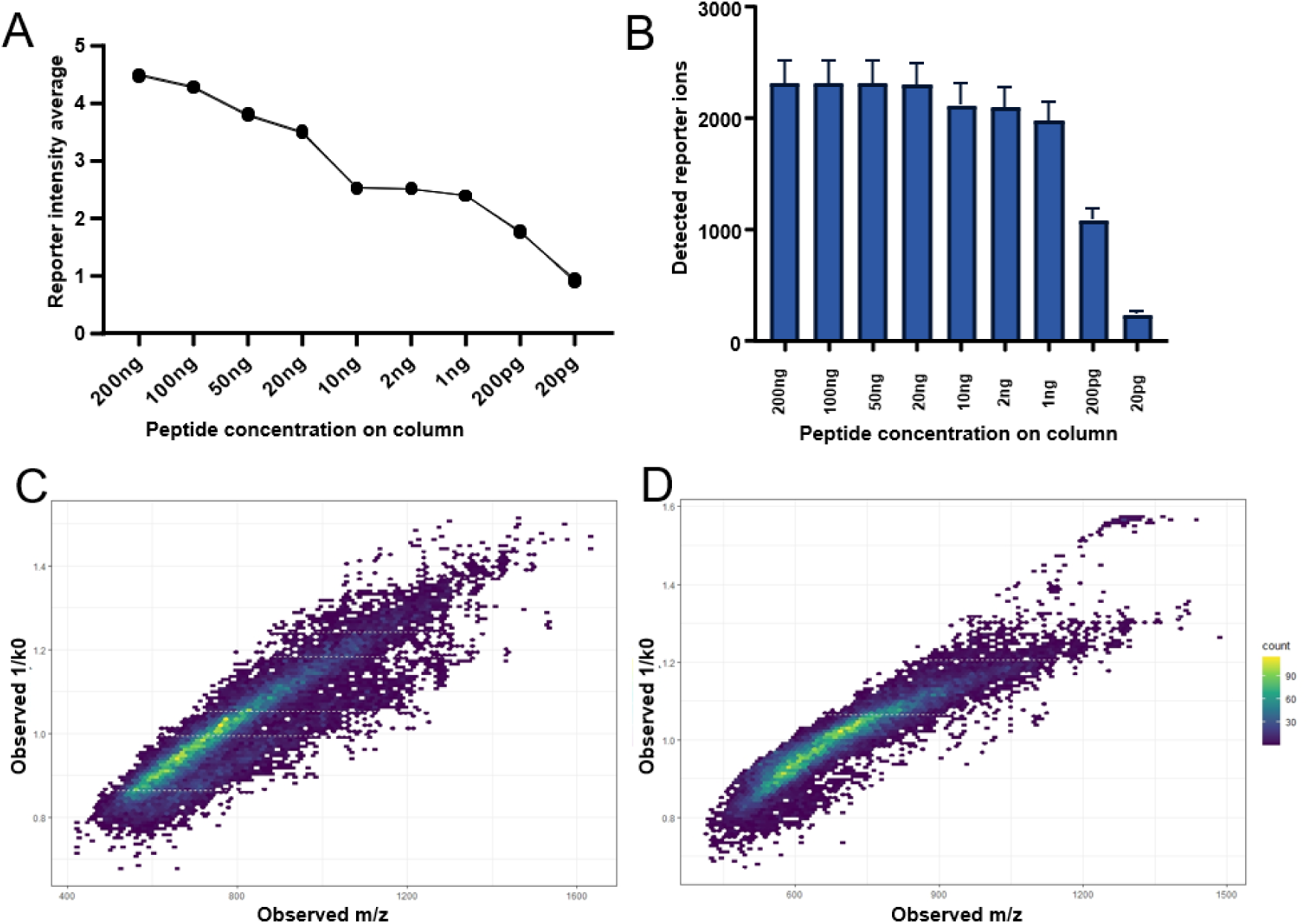
**A.** The log10 converted average intensity of each reporter ion in a TMTPro 9-plex linear dilution series. **B.** A comparison of the number of detected reporter ions at each concentration to evaluate the number of relative missing values across the dilution series. **C.** A plot of the distribution of m/z and 1/k0 values for unlabeled peptides. **D.** A plot of the same concentration of sample of peptides labeled with TMTPro reagent.

**Fig. 2.**
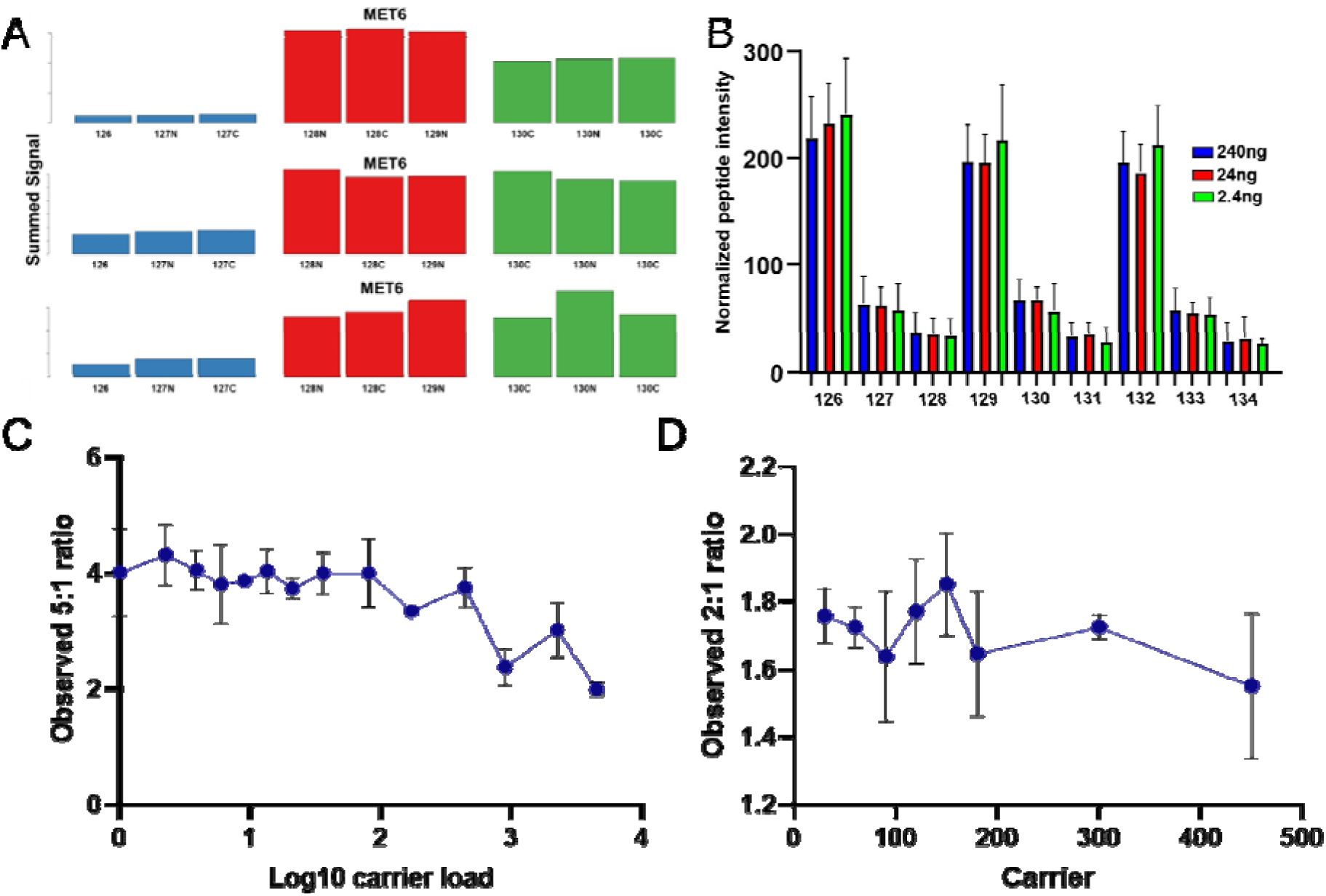
Assessing the background interference, quantitative accuracy, and carrier proteome effect of pasefRiQ. **A.** A comparison of results from 3 instruments that following analysis of 200ng of the TMT-TKO yeast digest standard and visualized using the TVT web tool.^28^ (Top) vendor example from an Orbitrap system using MS3 based quantification. (Middle) vendor example results from a quadrupole Orbitrap system using MS2 based quantification. (Bottom) Results from a pasefRiQ file. **B**. A graph demonstrating the normalized abundance of the *E. coli* protein TNAa in the two-proteome standard injected at different concentrations on column to illustrate the effects of sample dilution on ratio accuracy. **C.** Quantitative results of a standard protein with a known 5:1 ratio in a two-proteome standard digest with increasing amounts of carrier exceeding 4,000x carrier load. **D.** A summary of a second carrier experiment with more precise titrations across the upper limits reported for other hardware configurations.

### Ion mobility optimization reduces co-isolation interference

The unintentional co-isolation and fragmentation of background ions and their alteration of protein abundance measurements is a major challenge in multiplexed proteomics.^21^ To evaluate the level of background co-isolation interference in pasefRiQ we utilized a well-characterized yeast triple knock out TMT standard (TMT-TKO) designed for this purpose.^22^ In this standard, biological replicates of a parent strain and three separate transgenic strains with single genes removed are labeled with three TMT 11-plex labels. The TMT-TKO standard was analyzed with pasefRiQ using a consistent 60-minute LC gradient with adjustments made to the quadrupole isolation and TIMS ramp settings in each iteration of the respective methods. Reductions in the quadrupole isolation widths had minimal effects on the number of proteins and peptides identified per run, with the exception of a 1Da symmetrical isolation width which resulted in an approximate 18% loss in identified peptides and corresponding protein identifications. We therefore chose to use the minimum quadrupole isolation width that did not lead to a loss in peptide identifications, which was a 1.5 Da symmetrical isolation. (**S. Data. 1A**) In order to determine the appropriate ion mobility settings, we plotted the observed 1/k0 values and m/z for all identified peptides from a K562 peptide mixture both unlabeled and labeled with TMTPro reagent. As shown in **Fig. 1C** **& 1D** the majority of peptide signal was observed within a relatively narrow 1/k0 region. Peptides labeled with the TMTPro reagent exhibit a less symmetrical distribution than unlabeled peptides. By targeting the ion mobility range within the region of 0.8 -1.3 1/k0 we obtain the highest level of reduction in signal from the TKO channels, and therefore co-isolation interference for TMT labeled peptides.

In order to multiplex more than 10 samples with commercially available reagents today, isobaric reporter reagents with alternating N15 and C13 isotopes must be used. The neutron mass discrepancy in these two isotopes leads to a separation of m/z of approximately 0.006 amu. To fully resolve these reporter ions, current generation Orbitrap instruments are equipped with an optimized resolution of 45,000 at an m/z of 200. Orbitrap systems operating at this resolution obtain fewer MS/MS scans per experiment than typical label free experiments which obtain MS/MS scans at the much faster scans of approximately 15,000 resolution. The ability to multiplex up to 18 separate samples simultaneously is an attractive return on this loss in data acquisition rate.^23, 24^

During the calibration and tuning process, we can obtain estimates on the TIMSTOF Flex mass resolution that routinely achieves 40,000 at 1222 m/z. To determine the capacity of a TIMSTOF Flex to achieve higher multiplexing, we prepared commercially available human tryptic digests while using all 16 reagents following a manual calibration of the TOF resolution. As shown in **S.** **Fig. 2** we can obtain nearly complete baseline separation of the 127n and 127c reporter regions.

### Post-acquisition mass recalibration further reduces isolation interference

To evaluate the effects of the reporter ion mass accuracy on quantification we developed an offline manual calibration tool, the *pasefRiQCalibrator*, that writes a new spectral file following a manual calibration adjustment across a user- specified mass range. Following manual evaluation of multiple reporter ion masses, we can determine an appropriate adjustment factor for each file. Prior to manual calibration the most effective reporter ion isolation window for pasefRiQ files described in this study was approximately 50 ppm, falling within previously reported mass accuracy estimations for the TIMSTOF.^13^ Following manual adjustment, we can reprocess the same files using a 20 ppm mass tolerance window with no loss in reporter ions quantified, and a reduction in the mean signal intensity of the observed Δmet6 reporter ion of 56.4% (**S.** **Fig. 3**). By applying a calibration adjustment, we improve our ability to accurately extract reporter ion quantification even while using tighter integration tolerances (**S.** **Fig. 4**). Remarkably, despite the lack of complete baseline resolution of the reporter ion regions in the Δhis4 and Δura2 channels, we observe clear decreased abundance of these channels following recalibration (**S.** **Fig. 5**) While this was a promising development, the lack of baseline separation of reporter ions has been shown by others to result in the loss of quantitative accuracy in protein ratios with less extreme quantitative differences.^25^ For this reason, we chose not to perform higher multiplexed quantification with pasefRiQ with this iteration of the TIMSTOF hardware.

**Fig. 3.**
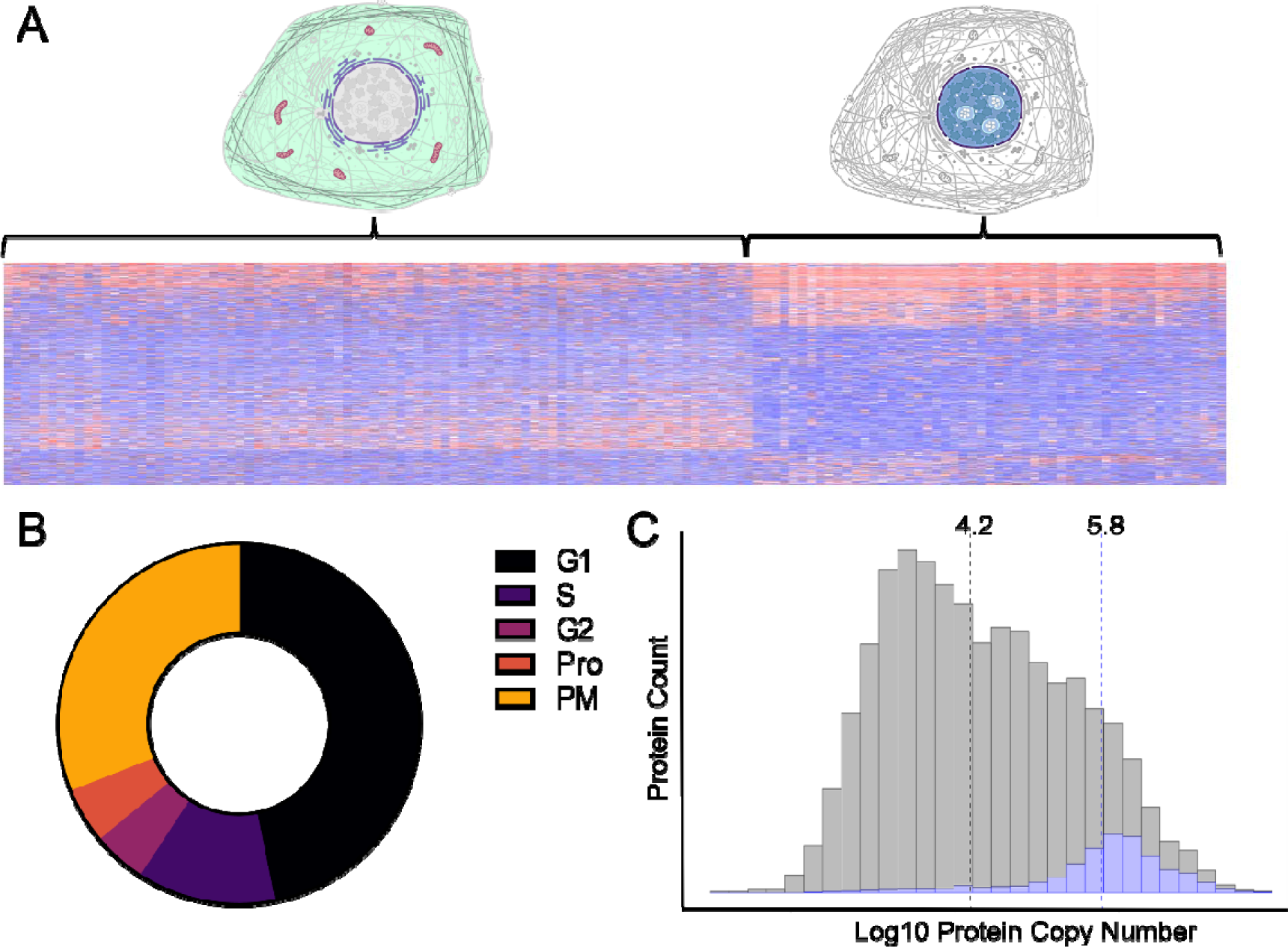
Characteristics of proteins quantified in single H358 cells. **A.** A heatmap demonstrating the main clustering characteristics of single cells. **B**. The cell cycle distribution of H358 cells as indicated by the relative abundance of 119 cell cycle protein models **C**. A histogram demonstrating the calculated copy number for 14,178 proteins (grey) compared to the proteins identified in single cells in this study (blue). Dotted lines indicate the median for each group.

**Fig. 4.**
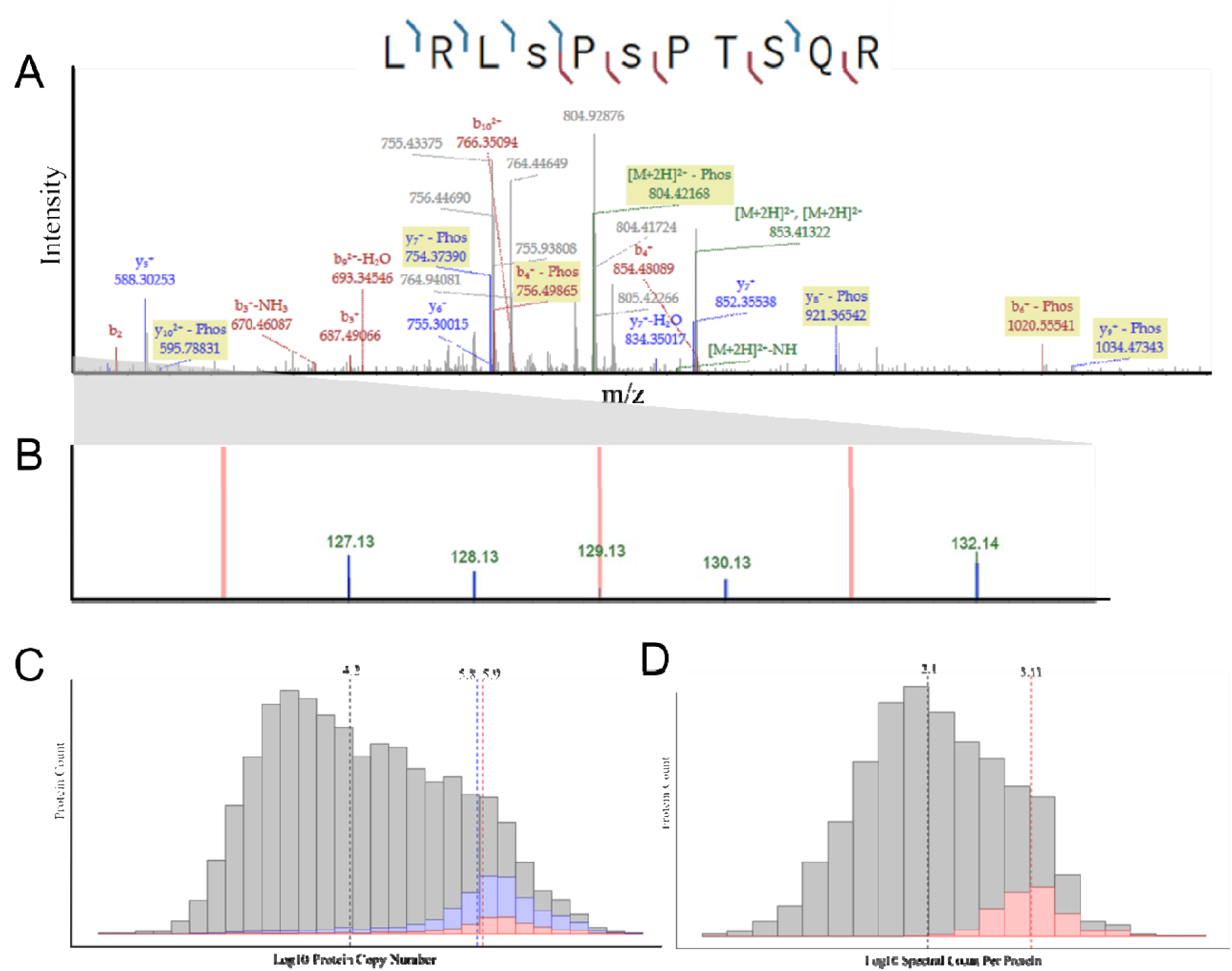
Phosphoproteins from single cells. **A.** A fragmentation pattern for LMNA phosphopeptide with sufficient fragmentation data for confident site localization. **B**. Zoomed in reporter region for this phosphopeptide demonstrating a lack of reporter signal in the method blank region and quantifiable signal from multiple single cells. **C.** A histogram with all proteins identified in single cells (blue) and phosphoproteins (red) compared against the calculated copy number of all proteins with median values highlighted. **D.** A histogram of the displaying the spectral counts for proteins identified in this study (grey) compared to the spectral counts of proteins where phosphopeptides were confidently identified. Dotted lines represent the median for each set of values.

**Fig. 5.**
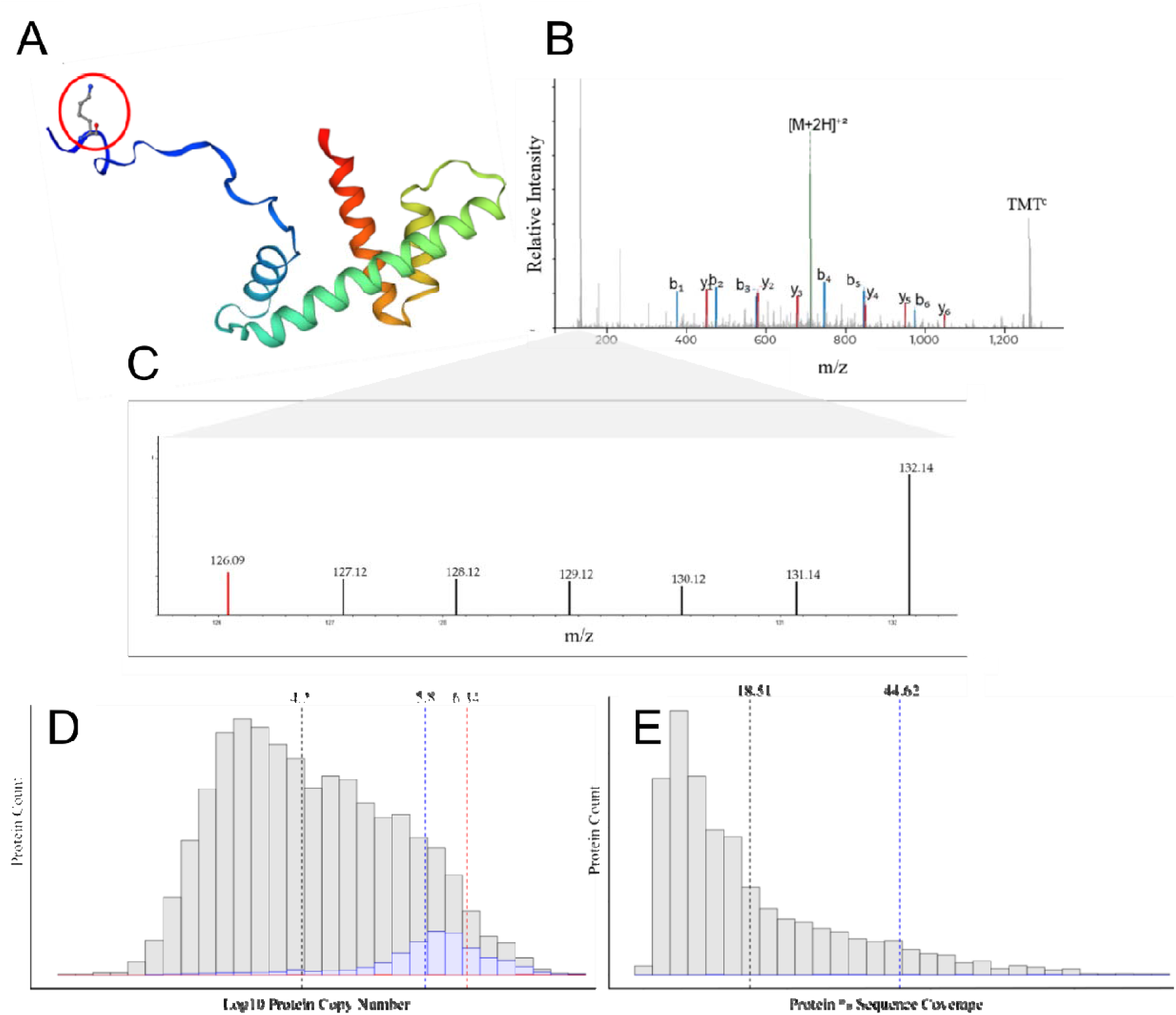
Acetylated proteins detected in single H358 cells. **A.** The Histone 2.2 protein with annotated acetylation site highlighted. **B.** A fragmentation spectra for this acetylation site demonstrating complete sequence coverage for identification and localization of this site. **C.** A zoomed in reporter ion region demonstrating signal for this peptide is apparent in every individual cell in this spectrum. The 126.09 highlighted in red is a diagnostic ion for an acetylated lysine residue. **D.** A histogram illustrating the relative copy numbers of all proteins in this study (grey), all proteins found in single cells (blue) and acetylated proteins (red). Dashed lines indicate median values. **E.** A histogram displaying a comparison between the total protein sequence coverage of all proteins identified in single cells (grey) and the median sequence coverage of acetylated proteins (blue line).

### Evaluation of quantitative accuracy with a two-proteome labeled 9-plex standard

The use of two-proteome standards is a well-established method in LCMS based proteomics for the evaluation of quantitative accuracy.^26, 27^ As such, we prepared 9 samples containing an identical concentration of a K562 commercial tryptic digest with a different level of *E. coli* tryptic digest spiked into each channel mixture to achieve a relative *E. coli* dilution series of 1:5:10 repeated three times within each LCMS run (**S. Data 1B**). To further evaluate the carrier load level between 100 and 500 x carrier which has been the focus of multiple studies using a single popular LCMS hardware configuration, an additional set of samples were prepared to provide a greater level of resolution within this dilution range (S. Data 1C).

### Quantitative accuracy of pasefRiQ is maintained at single cell relevant concentrations

To determine the relevant biological limits of detection of pasefRiQ we prepared serial dilutions of the two-proteome standards from 240 ng to 2.4 ng on column. While the number of peptides and proteins identified exclusively from MS/MS spectra at each subsequent dilution level decreased (**S. Data 1C**), we observed no decrease in quantitative accuracy. The expected ratio of the diluted *E. coli* proteins in this standard was 1:5:10 with 3 intraexperiment technical replicates across the 9 channels. When averaging the ratios from three separate LCMS experiments, the most accurate ratios for the TNAa protein observed were from the 2.4 ng injections which returned mean ratios of 1: 3.98: 7.43. The least accurate ratios were observed for 240 ng injections on column with mean ratios of 1: 3.28: 6.09 (**Fig. 2B**). These results demonstrate that values observed from pasefRiQ at picogram levels of peptide load per channel can return reliable quantification values.

### pasefRiQ is less restricted by carrier proteome effects

The effective amplification of reporter ions signal with carrier channels of increased relative concentration has limitations recently described as the carrier proteome effect.^19, 20^ To date, all analysis of this effect have been performed on various iterations of hybrid Orbitrap architecture. To assess this effect in pasefRiQ we again employed the two proteome standard digest model supplemented with increasing amounts of peptides labeled with the 134n channel. As shown in **Fig. 2C**. for a protein with a known relative ratio of 5:1 between labeled channels we observe no meaningful change in this ratio when employing a carrier channel of up to 500-fold higher concentration than the other respective peptides. In addition, when employing a carrier channel in excess of 4,000-fold beyond that of any respective channel, the known 5:1 ratio, while compressed, was still observed as a 2:1 ratio (**S. Data 2**). A second carrier dilution range inspired by the multiple studies conducted on the popular D20 Orbitrap instruments^6, 19, 20^ was prepared to provide greater resolution across the upper limits where these instruments have been demonstrated to suffer quantitative effects. We again see less relative distortion than previously reported (**Fig. 2D**). As described by others, we do observe quantitative impurities at high carrier loads.^20^ When employing a carrier channel of approximately 500-fold higher concentration relative to all other channels as label 134n we find inflation of the protein summed intensity in channels 133n and 132n (**S. Fig. 7**).

### Over 1,000 proteins can be identified in single cells in 30 minutes

To evaluate the capabilities of pasefRiQ for the analysis of single human cancer cells, individual NCI-H-358 (H358) cells were aliquoted by flow-based sorting into 96-well plates. The first well in each row was loaded with buffer with no cell to provide a channel for intraexperiment estimation of background interference. To determine the optimal carrier channel concentrations, a dilution range of peptides from an H358 bulk cell lysate was tested as potential carriers to determine the ideal concentration for protein detection without suppression of single cell signal (**S.Fig. 8A**). A bulk cell lysate of H358 cells labeled with channel 134 at an approximate concentration of 50 ng or 250x, that of a single cell was selected from these data as the optimal carrier channel concentration. (**S.Fig. 8B**). To maintain a high daily throughput of single cells with pasefRiQ reported techniques we chose to use 30-minute gradients which were approximately one third the total LCMS acquisition time of previously reported studies (**Table 1**).

**Table 1.** A comparison of pasefRiQ single cell results to Orbitrap MS2 based publicly available data

| Study | # MS2 scans | #Peptides | #Proteins per cell | Max # of cells | Time |
| --- | --- | --- | --- | --- | --- |
| This study | 42555 | 8515 | 1044 | 6 | 30 min |
| SCOPE2 | 8418 | 2450 | 599 | 9 | 100 min |
| nPOP | 8248 | 1822 | 775 | 14 | 105 min |

To determine the relative level of performance of pasefRiQ for single cell proteomics, a plate of H358 cells was prepared and analyzed within a single batch (**S. Fig 9**). The resulting pasefRiQ files were analyzed alongside previously published single cell data using the same software, settings and quality filters. These results (**Table 1** and **S. Data. 2**) demonstrated comparable numbers of protein identifications could be obtained by pasefRiQ in approximately one-third of the total instrument acquisition time. In addition, we observe an approximate five times more fragmentation spectra per experiment and a corresponding increase in the number of peptides and an overall increase in total sequence coverage for all identified proteins (**S. Fig. 10**)

### Analysis of 443 single cells in 50 hours of total instrument time

To further expand on this study, proteomic data was acquired on a total of 443 single H358 cells using approximately 50 hours of LCMS instrument time. This time was inclusive of control runs and approximately 15 minutes of equilibration between experiments required by our system.

The spectral data produced by TIMSTOF instruments is unique in many ways when compared to any previously generated mass spectrometry data. As such, only a relatively small number of historic data processing pipelines are currently compatible with these data, and all that are currently have limitations of some type (**S. Table 1**). To obtain the most comprehensive interpretation of peptide and protein identifications we utilized four search tools in tandem. In total, 2,125 proteins were identified, with 1,858 proteins identified by at least two search tools. The most conservative search tool, MaxQuant, identified 1,631 protein groups at an average of 655.2 quantifiable proteins per cell, with a maximum of 1,255 proteins with reporter ions corresponding to a single cell. MSFragger had the highest identification rate with 1,813 protein groups identified with an average of 746.2 per single cell. When combining the results of all proteins identified by all four tools, we obtained an average of 946.3 proteins per cell. (**S. Fig. 11; S. Data 3, S Data 4**.) The number of proteins identified in this cell line was comparable to previously described iterations of the SCoPE-MS workflow reported by others when using cancer cell lines for analysis.^6^

### The cell cycle status of each individual cell can be estimated using proteomic markers

A central tool in scSeq data analysis relies on the removal, grouping or flagging of the markers of individual cell cycle status to keep these large effects from compromising other analyses.^29, 30^ To date, no parallel approach has been described for single cell proteomics. As described by others, unsupervised clustering of the proteomes of individual H358 cells are largely driven by the cell cycle status of each individual cell, suggesting the possibility and importance of cell cycle flagging techniques.^31^ When using a global unbiased approach we observed the majority of cells clustered into two main groups differentiated by the relative abundance of proteins linked to metabolism, translation and cell division (**Fig. 3A**). When using the detection and relative abundance of 119 protein cell cycle markers for guidance ^32^ we find that we can estimate the cell cycle status of each individual cell. Using this approach we determined that the majority of cell analyzed demonstrated prominent G1 and prometaphase markers, respectively with the remainder of cells demonstrating markers consistent with cell cycle status between these two (**Fig. 3B** and **S**. **Data 5**). We have provided a simple curated protein database compatible with any LCMS proteomics workflow for the estimation of the cell cycle status based on these protein markers as well as graphical tools compatible with the software used in this study along in the repositories for this manuscript.

### Proteins quantifiable in single cells track closely to protein copy number estimates

We have previously described a public Shiny Web application for the estimation and filtering of proteomics data against libraries of established cellular protein copy numbers.^33^ Using this tool we find that proteins identified in H358 single cells closely track to the theoretical copy number for each individual protein. Over 95% of all proteins with quantifiable signal from individual cells in this study were from proteins estimated to possess more than 100,000 copies per cell. The copy number for a protein identified in a single cell in this study is approximately 630,000 copies. In contrast, the median calculated copy number of all proteins in this cell line is approximately 16,000 copies (**Fig. 3C**).

### Protein post-translational modifications are confidently identified in single human cells

To date, no single cell study using mass spectrometry has described the identification of protein post-translational modifications (PTMs). To evaluate if the increase in relative sequencing information in pasefRiQ could provide insight into PTMs, pasefRiQ single cell files were analyzed using well-established PTM identification pipelines.^3^ Over 2,000 high confidence peptide spectral matches were made for sequences containing phosphorylation, acetylation, methylation, dimethylation, succinylation, hydroxybutylation, crotonylation and cysteine trioxidation (**Table 2, S. Data 6**). Many abundant PTMs were identified with supporting evidence across multiple LCMS runs and with reporter ions corresponding to all single cells passing our quality control pipeline checks. Searching for PTMs expands the search space and leads to an inflation in potential false discoveries. When comparing the peptide spectral matches made when searching for these PTM using our pipeline to a search made without PTMs, only 23 MS/MS scans (0.014%) were assigned to an alternative identification (**S. Data 7**). These results indicate that our search parameters produced no meaningful inflation in false discovery rates. For further confidence in identifications, all PTMs reported herein have been manually examined for sequence quality. All peptide spectral matches reported for PTMs in this study have been made available through a web interface featuring visualization tools for matches and decoy match identifications.

**Table 2.** A summary of PTMs identified in single cells in this study.

| Modification | Modified proteins observed | Cells where this PTM was observed (%) | Most observed modification site | Cells where this site was observed (%) | Previously characterized PTM |
| --- | --- | --- | --- | --- | --- |
| Acetylation | 32 | 95.7 | Histone H3-3A K24 | 49.1 | Yes |
| Phospho | 79 | 100.0 | NUCKS1 S181 | 43.9 | Yes |
| Methylation | 20 | 100.0 | Histone H3-3 K80 | 54.3 | Yes |
| Succinyl | 6 | 66.1 | GNG5 K11 | 52.2 | No |
| Dimethylation | 23 | 100.0 | Histone H3-4 K28 | 54.8 | Yes |
| Hydroxybutyryl | 1 | 0.9 | WASH-2 K661 | 0.9 | No |
| Crotonyl | 10 | 90.0 | RPL14 K164 | 67.4 | No |

### Characterization of phosphoproteins identified in single cells

The most common phosphorylated protein observed in single cells was the nuclear prelaminin protein A (LMNA), which was identified as phosphorylated in 153 of the cells analyzed in the total course of the study. The most common sites observed on the protein are well-characterized serine sites Ser22, Ser390, Ser392, Ser404, and Ser406 that were observed in 114 individual cells in total with other phosphorylation sites on this highly modified protein observed in additional cells. Sufficient fragmentation data was obtained for localization of the site of phosphorylation in peptides where multiple potential sites were observed.^34^ (**Fig. 4A/4B**). The abundance of LMNA phosphopeptides allows for clear reporter ion signal to be observed for multiple single cells within a single spectrum (**Fig. 4B**). To add further confidence to these identifications, all files were independently analyzed with the MSFragger^35^, MSAmanda^36^, and Sequest^37^ search engines using search parameters as similar as possible within the limitations of each user interface. The output results of these files were combined and judged independently by semi-supervised machine learning using the Percolator^38^ and phosphoRS^34^ algorithms to obtain consensus results. In total, 889 phosphorylation spectra were identified from all proteins at less than 1% false discovery rate (FDR). These peptide spectral matches condensed to 89 unique high confidence phosphorylation sites occurring in total on 37 separate proteins, with a high relative agreement between all search tools. Of the 114 phosphorylation sites on LMNA observed in single cells 85/114 (74.6%) were confirmed by all three search engines, and all 114 were confirmed by at least two tools (**S. Data 8**).

When compared to the calculated protein copy number, phosphoproteins were found to have a median abundance of approximately 600,000 copies (**Fig. 4C**), a value only 25% higher than the median for the identification of proteins themselves in single cells. However, a clear difference was found when comparing the amount of sequence information obtained for each protein from the two groups. Phosphopeptides were identified on proteins containing over 10-fold supporting peptide spectral matches than identified proteins as a whole (**Fig. 4D**).

The relationship between protein copy number, spectral counts and confident phosphorylation site location was highlighted by the multiple sites observed on PLEC and AHNAK. Over 1.9% of all MS/MS spectra acquired in this study (51,075/2,599,602) can be attributed to the 531 kDa PLEC cytoskeletal protein. Similarly, the 628.7 kDa AHNAK protein comprised 1.3% of all total MS/MS spectra in this study. The 34,311 PSMs allow over 76% sequence coverage of this protein with quantifiable signal observed in nearly every cell in the study. At the protein sequence level, proteins with a single phosphorylation site had a median total sequence coverage of 39.4%, compared to the average protein which contained 16.1% sequence coverage (**S. Data 8**).

### Mitotic phosphopeptide abundance correlates with the expression of other mitotic proteins

Phosphorylations on nuclear laminin protein LMNA have well characterized functions in mitotic regulation. Phosphorylation of Ser22 and Ser392 were detected in 33 and 22 individual cells, respectively. These “mitotic sites” are essential for LMNA localization and actively promote depolymerization of the intact nuclear lamina to allow nuclear division.^39^ In contrast, phosphomimetic mutations phosphorylation of Ser390 demonstrated no observable alterations in nuclear location of LMNA.^40^

Proteins quantified in single cells in this study with annotated involvement in specific cell cycle stages and their relative abundances were extracted from the whole proteome processed reports. Pearson correlation coefficients were calculated using the relative abundance of each LMNA phosphopeptide in each cell against the abundance of each cell cycle protein (**S. Fig. 13A** and **S**. **Data. 9**). Both Ser22 and Ser392 phosphorylation showed a strong correlation to multiple proteins involved in the G2/M transition. In addition, Ser392 phosphorylation was also strongly correlated to later stages of mitosis than Ser22. Ser390 phosphorylation showed no correlation to mitosis related protein abundance except for strong negative correlation to TUBB4B. Ser390 phosphorylation only demonstrated strong correlation to a single mitosis protein, the regulator of ploidy, LATS1 (r =0.9980) with a weaker correlation to and RAB8A (r = 0.78).

When extending this correlation analysis of these three phosphorylation sites to all proteins quantified in H358 single cells, we again found that Ser22 and Ser392 demonstrated the strongest correlations to proteins annotated by gene ontology as cell cycle mediated. Ser390 phosphorylation most strongly correlated with EIF2AK2 and PML, which are known to be involved in the inactivation of protein translation. (**S. Fig 13B**).

These results lend support to both our PTM identifications and to the value that the quantification of PTMs may play in elucidating the intracellular environment in single cells. While considerable work has been performed using molecular biology approaches to the study of LMNA phosphorylation, we can find no published accounts detailing the interplay of these phosphorylation sites at the single cell level (**S. Fig 13C**). In addition, it may be possible through single cell proteomics to assign putative functions to phosphorylation sites such as LMNA Ser390 through the application of single cell proteomics.

### Characterization of modified histone proteins

Lysine acetylation is a PTM with regulatory importance in a variety of cellular systems.33 Notably, lysine acetylation plays a central role in the regulation of nuclear histone proteins.34 We observed 32 high confidence acetylation sites in single cells, primarily localized on well-characterized acetylation sites within histone proteins (**Fig. 5B, S. Fig 12)** is an example of one such site from Histone 2.2. Peptides with an acetylated lysine commonly produce a diagnostic fragment ion during collision induced dissociation with a mass of 126.0913. Mass spectrometers with lower relative resolution or mass accuracy may not be able to accurately discern the 0.0364, or 29ppm mass difference between this diagnostic ion and the 126 TMT reporter ion. As shown in **Fig. 5C** we can confidently extract reporter ion signal from each H358 cell analyzed within that spectrum and observe no reporter ion signal from the 126 method blank control well, while clearly discerning the lysine acetylation diagnostic ion. Diagnostic ion filtering of spectra identified 35,130 spectra, or 3.6% of all filtered spectra contain a putative lysine acetylation diagnostic ion. When compared to the number of spectra identified for peptides from a single histone such as 2.2 which was supported by over 2,200 separate MS/MS spectra in this study, these results are not altogether surprising. Histone proteins are among the most abundant within mammalian systems, often occurring in excess of one million copier per cell. This abundance has a direct effect on the characteristics of proteins where acetylation sites were observed. Acetylation sites were identified on proteins with a median log copy number of 6.38, or approximately 2.2 x 10^6^ copies per cell, representing proteins in the top 3% of total predicted abundance. The value of total protein sequence coverage was also apparent in the identification of acetylated proteins in single cells. The median total protein sequence coverage for a protein with a confidently identified acetylation site is 44.62%, compared to a median of 18.51% for all proteins identified in this study.

The occurrence of protein methylation and dimethylation sites follows a nearly identical pattern to acetylation in that they were almost entirely detected on high abundance histone proteins such as H3-3 and H3-4, respectively. The Histone H3-4 dimethylation site localized to lysine 28 was supported by 878 MS/MS spectra and reporter ions from this single modification can be confidently assigned to 54.8% of all single cells in this study. Histone proteins are challenging to identify despite their high abundance as they are both heavily modified and extremely basic. In order to obtain high sequence coverage of most histones derivatization of unmodified lysines and other strategies are often used to increase the length of the average peptide sequences observed by LCMS.^41^ These results suggest that derivatization methods compatible with single cell proteomics processing may provide deeper insight into the epigenetic landscape of single cells.

### Additional PTMs identified in single cells

Lysine crotonylation is PTM of recently revealed importance in a variety of cellular mechanisms, including DNA damage repair and carcinogenesis.^42^ Ten unique crotonylation sites were observed on single cells in this study. Although crotonylation is typically associated with histones, none of the identified sites were confirmed on histones in single cells. The most observed crotonylation site was on 60s ribosomal protein L14 (RPL14), which was confidently identified in 67.4% of all cells in this study. Recent studies have implicated protein crotonylation in the activity of cells with KRAS mutations. When comparing two human non-small cell lung carcinoma cell lines, 7,765 crotonylated peptides were identified in A549, a KRAS^G12S^ mutant cell line, nearly 3- fold more than were observed in NCI-H1299, a KRAS wild-type cell line.^43^ Of the crotonylation sites observed in the A549 KRAS mutant line, 346 correspond to modifications on 60s ribosomal proteins, including RPL14.^44^

Although 52 peptides were identified with a putative cysteine trioxidation with signal corresponding to single cells, manual analysis of these identifications did not provide sufficient sequence coverage to adequately support these identifications. Finally, a single high confidence hydroxybutyrylation site was identified in single cells in this study, which was localized to the K661 residue of the actin regulating protein WASH-2. Although acyl based modifications have been implicated in the function of WASH-2, this site has not previously been characterized and was only identified in four single cells in this study.^48^

### Application of single cell proteomics to a drug treatment model

To explore the power of single cell proteomics toward drug mechanism studies, we treated H358 cells with the FDA approved KRAS^G12C^ covalent inhibitor, sotorasib, using the same culture and dose concentrations described in a recent single cell RNA-seq (scSeq) study of the same.^49^ Following data filtering and normalization (**S. Fig 14**), the effects of drug treatment can be clearly discerned by simple tools such as principal component analysis (PCA) (**Fig. 6A**). When cells treated with sotorasib were analyzed as if they were technical replicates with peptide and protein abundances averaged, our results closely mimic the effects of this compound as established by others (**S. Fig. 15A**). Gene set enrichment analysis (GSEA) of proteomic alteration found the canonical VEGF pathway to be the single most altered mechanism upon sotorasib treatment (**S. Fig. 15B**), in line with previous observations.^50^

**Fig. 6.**
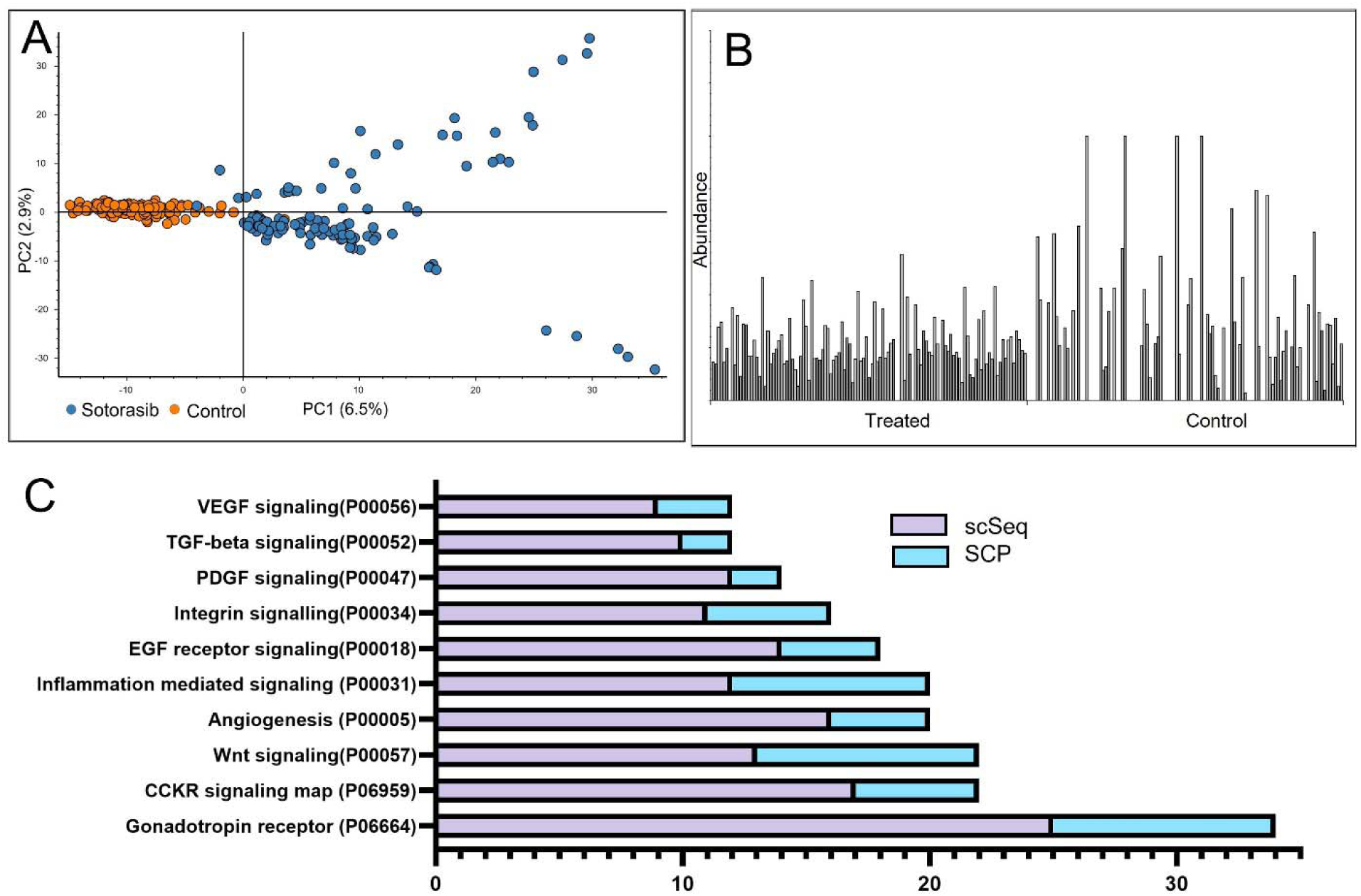
Single cell proteomics provides insight into cellular response to drug treatment. **A**. A PCA plot demonstrating PC1 (6.5%) and PC2 (2.9%) of proteins from single control and sotorasib treated cells. **B.** The measured abundance of TOP2A protein expression in 230 single cells. **C.** The top pathways identified as differential by a scSeq analysis of drug treatment combined demonstrates the strengthening of each pathway when SCP data is added.

When compared to the results of a scSeq analysis of H358 cells treated with the same dose of inhibitor we find that 12.1% of protein identifications are decreased by more than 2-fold directly overlap with the transcript data filtered at the same level. However, when examining proteins and transcripts identified as differential in a functional context, we find the data to be even more complementary as nearly every pathway identified as differential was enhanced when the two datasets were combined (**Fig. 6C, S. Data 10**) In addition, scSeq analysis of sotorasib treatment identified alterations in the cell cycle distribution of single cells versus control. Following treatment we observe similar changes in the protein markers of cell cycle, highlighted by a 4-fold increase in prophase proteins in drug treated cells relative to control (**S. Fig. 16**).

### Applying single cell proteomics to assess the level of heterogeneity in drug response

A recent temporal proteomic analysis of two cell lines treated with KRAS^G12C^ covalent inhibitors identified proteins and phosphopeptides with differential expression following treatment.^51^ When treating single cells as replicates, we find our data to be in generally high concordance with these observations, despite the lower number of quantified proteins in single cells. To evaluate the relative consistency of proteomic response across the cellular population we utilized tools capable of visually representing the relative expression of proteins and transcripts across hundreds of cells simultaneously.

For example, the proteomic analysis of cellular lysates treated with sotorasib found a relative increase in abundance of the DNA damage response protein, Death Associated Protein 1 (DAP1) of 2.23-fold. When evaluating the expression of DAP1 in single cells treated with inhibitor we find a DAP1 fold change at the 100-fold maximum ratio allowed by our quantitative tools. When visually inspecting the output from single cells we find that DAP1 expression was detected exclusively in 43/115 treated cells, where it was never detected in a single control cell (0/115) (**S Fig. 17A**). When examining DAP1 transcript expression at the single cell level, we find a moderate increase in transcript levels with treatment at 24 and 72 hours, however, the raw transcript abundance of DAP1 places it near the lower limits of detection in the single cell data.^49^ These results suggest that DAP1 protein expression is increased by sotorasib treatment, but this effect, while common, may not be true for an entire cellular population.

In a second example TOP2A is a protein that was detected in 81.3% of all single cells in this study. The relative 3-fold decrease in TOP2A abundance following drug treatment appears to be a mostly homogenous response that closely mimics the observed 3-fold decrease in the bulk cell homogenate data (**S. Fig. 17B**). These results are an almost exact match in the scSeq data, when averaging the relative expression of approximately 1,000 single cells where a decreased abundance of 3.33-fold was observed following sotorasib treatment (**S. Fig. 18**).

One contrary example was observed in the relative expression of the chloride channel protein CLIC3. In both the bulk temporal proteomics and single cells when protein expression is summed, we observe marked differential abundance of CLIC3. Closer evaluation at the single cell level found that this differential in abundance was driven by ten treated cells that demonstrate a high level of CLIC3 expression, while no quantifiable signal was observed for this protein in any other cell (**S. Fig. 19**). These results suggest that the increased abundance of this chloride channel protein may be an extreme phenotypic response of a relatively small cellular population at this timepoint. To evaluate the potential differences leading to the CLIC3 expression phenotype, these cells were analyzed as a separate subgroup from all other sotorasib treated cells. StringDB pathway analysis strongly implicated translation initiation and protein translocation to the endoplasmic reticulum were the most enriched functional pathways of this small cellular subgroup. (**S.Data.11**) While this may suggest that the CLIC3 phenotype is an marker of cells that have overcome the sotorasib induced quiescence phenotype, further study would be necessary to make this conclusion.

## Conclusions

Single cell proteomics is a newly emerging field with tremendous promise for new biological discoveries. Single cell studies described to date have necessarily focused on developing the sample preparation,^6, 10, 52–55^ instrument methods ^7, 31^ and informatics tools ^56,57^necessary to lay the groundwork for later applications. Much of this valuable development has been performed using diluted bulk proteomics samples ^19, 58^, large cell types from other model organisms ^59^, or clonal populations of hundreds of cells harvested from single cell seeding or laser capture microdissection.^60^ Many of these studies have contributed key insights into the challenges LCMS faces at picogram concentrations of peptide. From both previous results and the protein identifications made in this study, it appears the primarily limiting factor in single cell proteomics is primarily the total concentration of each protein in a single cell, despite the hardware configuration utilized.

In this study we have detailed optimized methods for reporter-based quantification with a focus on reducing background coisolation interference and obtaining the highest quantitative accuracy. We find that single cell proteomics by pasefRiQ can provide accurate quantitative information at single cell-relevant concentrations and is less hindered by the “carrier proteome effect” than other hardware configurations. Due to the increased relative speed of data acquisition of the TIMSTOF instruments we can obtain dramatically more sequence information for each protein identified and this information allows the identification of protein post-translational modifications. As demonstrated by others, we find that cell cycle linked proteins and their abundance impact the proteomes of the cells observed and that this extends to cell cycle regulated PTMs which exhibit a strong correlation to the appropriate cell cycle markers. We have developed simple tools that can be used for estimating the cell cycle status of each individual cell and allow these effects to be removed from confounding other biological interpretations. The high relative intracellular abundance of histone proteins allows for the confident identification of histone acetylation, methylation and dimethylation sites in the majority of cells analyzed. While further work is clearly necessary to build the informatics framework and methods toward the identification of other PTMs such as protein glycosylations, preliminary evidence presented here suggests that single cell glycoproteomics may be a promising future application.

Finally, we present the application of single cell proteomics to a drug mechanism study of single cells by treating a model KRAS^G12C^ mutant cell line with the covalent inhibitor sotorasib. Sotorasib was approved by the FDA in late 2021 as the first in a line of similar covalent inhibitors currently in clinical trials and under development.^14^ While possessing clear value as a first of its kind treatment, spontaneous resistance to this drug has been observed in both cell lines and patients.^61^ Recent work using temporal proteomics identified multiple proteins upregulated following inhibitor treatment and demonstrated the value of combinatorial treatments that inhibited both KRAS^G12C^ and these resistance mechanisms.^51^ When comparing our results to the temporal bulk proteomics and other related studies of KRAS^G12C^ inhibition, we find our results generally high concordance. We observe both suppression of KRAS itself, as well as a number of proteins in the overlapping VEGF and MAPK pathways that are hyperactivated by the perpetually GTP bound and active KRAS mutant protein.^62^ When considering our cells individually, we find these results less clear and more reflective of the heterogeneity in response identified by a scSeq study of this drug. While proteins in the MAPK pathway appear clearly downregulated, with most observable error linked to the stochastic nature of this method, some protein level observations appear to be wholly driven by extreme protein upregulation in small cell populations. Furthermore, we find that LMNA phosphorylations observed in the bulk cell proteomics were likely confounded by cell cycle mediated effects where these phosphorylations play a key role. Cell cycle status was recently implicated as a key driver in the development of sotorasib resistance, suggesting that further investigation into these mechanisms through single cell proteomics would be a valuable contribution to the understanding of inhibitor response and resistance. We find these results highly promising as indicative of the power single cell proteomics can play in pharmacology studies.

## Materials and Methods

### Samples for optimization

Human cancer cell line digest standards K562 (Promega) and diluted to 100 microgram/mL in 50mM TEAB and labeled with TMTPro reagents according to manufacturer instructions. For preparation of the TMTPro 16 and TMT9 standards, the unit resolution reagents 126, 134, 127n, 128n, 129n, 130n, 131n, 132n, and 133n channels were used in all experiments. The loading capacity of the LCMS system was determined empirically from dilution series of these standards. A TMT triple knock out (TMT TKO) yeast standard (Pierce) was prepared following manufacturer instructions and prepared in serial dilutions in 0.1% trifluoroacetic acid.14 The two proteome standard for assessing quantitative accuracy was prepared from K562 digest as above with an *E.coli* peptide standard digest (Waters MassPrep 186003196). The standards were mixed and labeled with the TMT9-plex standard to maintain a constant concentration of K562 digest in all 9 channels while the *E.coli* peptides were prepared in three separate concentrations in triplicate to maintain a known ratio of 1:5:10 of E.coli peptides in triplicate across each 9-plex. For carrier proteome analysis two additional samples were prepared where the 126 and 134 channels, respectively, were substituted with a 1:1 ratio of K562 to *E.coli* a mixture of the other 8 channels was diluted and combined with the carrier mixture to approximate carrier loads of 1x, 2.25x, 3.9x, 6x, 9x, 13.5x, 21x, 36x, 81x, 161x,171x, 441x, 891x, 2241x and 4491x relative to the concentration peptides in each individual sample To further titrate the region of the carrier proteome where previous studies have found quantitative ratio distortions, a second sample set was prepared using the 135n channel as carrier.^6, 20, 58^ In this sample set, *E.coli* dilutions in 126- 129 were 1:2:5:10 and repeated in 130-133. Five carrier loads were used as 135n, at 30x, 60x, 90x, 120x, and 180x, respectively

### LC Orbitrap based analysis of 2-proteome standard for internal comparisons

For comparative analysis of the two proteome standard a sample containing a total peptide load of 240ng was analyzed on a Q Exactive “Classic” mass spectrometer with an EasyNLC 1200 system and utilizing a 15cm C-18 PepMap100, 2 µm EasySpray column (150 mm x 75 µm, ES904) and an Acclaim PepMap 20 mm trap column containing the same chromatography material (164946) (all components from Thermo Fisher). Peptide separation was performed using a gradient with a total length of 86 minutes with a separation ramp of 8% buffer A (0.1% formic acid in LCMS grade water) to 24% buffer B (0.1% formic acid in 80% acetonitrile) in 60 minutes, followed by an increase to 36% B by 70minutes, followed by a rapid ramp to 98% B which was held for the remainder of the run. The Q Exactive was operated using a previously curated method deposited in the LCMSMethods.org 2019 method’s collection title QE_Plus_iTraq4_TMT6.meth. Full instrument methods and details have been published (https://dx.doi.org/10.17504/protocols.io.bzd4p28w). Briefly, MS1 spectra were acquired at 70,000 resolution from 400-1600 m/z with a maximum AGC target of 3e6. With peptide match employed as “best” the top ten ions from each parent scan were isolated at 1.4 Th using with an ion target of 2e5 and maximum fill time of 114 ms with 17,500 resolution. Isolated ions were fragmented with a normalized collision energy of 28. Due to the asymmetrical quadrupole isolation of the single segment quadrupoles previously reported, a lower isolation window was not performed on this instrument.^63^ Singly charged ions, those with more than 7 charges or those with undetermined charge state were excluded from fragmentation. Ions selected for fragmentation were excluded from additional fragmentation for 90 seconds using a mass tolerance window of 5ppm from the isolated parent. Isotopes of selected ions were likewise excluded. For comparative analysis, Orbitrap and pasefRiQ analysis of 240ng peptide load were processed within a single workflow in Proteome Discoverer 2.4 and comparisons were performed using the 1083 overlapping proteins quantified in both instances.

### Preparing bulk samples for spectral libraries and carrier channels

NCI-H-358 cells were obtained from ATCC (Catalog #5807) and were reconstituted and passaged according to the included instructions using 6 well culture dishes. For bulk cell experiments, cells were aspirated, washed with ice cold water which was rapidly aspirated prior to addition of S-Trap lysis buffer. All steps of the S-Trap mini protocol were performed according to manufacturer instructions (ProtiFi) with the exception that alkylation and reduction were not performed. Peptides for spectral library generation were labeled with the 128C reagent from the TMTPro reagents according to all manufacturer protocols. The peptides were fractionated by using high pH reversed phase spin columns (Pierce) and eluted peptides were centrifuged to near dryness prior by SpeedVac. Peptides for use as carrier channels were labeled with the 134N reagent from the TMTPro, lyophilized and the concentration was determined using a Qubit system (Thermo Fisher) following manufacturer instructions.

### Preparation of single cells

Single control cells were treated with DMSO or with a 10 µm sotorasib solution (SelecChem S8830) prepared in DMSO as previously described.^49^ Sotorasib treated, and control cells were cultured alongside control cells for 40 hours prior to rapid washing of cells with ice cold magnesium and calcium free PBS (Fisher). Cells were sorted and aliquoted following a previously published protocol using flow cytometry. Briefly, cells trypsinized to circularize, washed and resuspended with trypsin inhibitor and rapidly taken to the Johns Hopkins University School of Public Health Cell Sorting and Sequencing Core on ice. Cells were stained with a viability marker and sorted directly onto microwell plates and were immediately transferred to dry ice prior to -80C storage.

Lysis, digestion, and labeling followed the SCoPE2 protocol previously described with minor alterations. Briefly, single cells were removed from -80C storage and placed directly in a solid fitted heat block at 95C for 10 minutes. (Fisher) All manipulation of plates was performed with the author grounded by alligator clip to the bench surface to prevent static discharge removing the cells from the microwells. The protein from lysed cells were digested in 1 µL 10ng/µL of trypsin/LysC (Promega) in 100mM TEAB (ProtiFi). Digestion was performed at 37□C for 3 hours in sealed plates on a revolving incubator. Plates were centrifuged at 4,000 x g at 4C every half hour to concentrate condensate. Following digestion, cells were labeled with previously aliquoted and stored TMTPro reagent 63 resuspended in LCMS grade anhydrous acetonitrile (Fisher) to a total concentration of 44mM. 500nL of resuspended reagent was added to each cell as appropriate and labeling was performed at room temperature for 1 hour. Plates were centrifuged twice to concentrate condensate. TMT labeling was quenched by the addition of 500nL of 0.5% hydroxylamine and centrifugal shaking for 1 hour at room temperature. Well with single cells were resuspended with the serial addition of the TMT134N carrier channel at an approximate concentration of 50 ng.

### LCMS settings for standards

All instrument settings are included within the Bruker. d files in the ProteomeXchange and have been uploaded to www.LCMSMethods.org as pasefRiQf_v1 and published as (dx.doi.org/10.17504/protocols.io.b4ifqubn). **Table 3** is a brief summary of these settings.

**Table 3.**
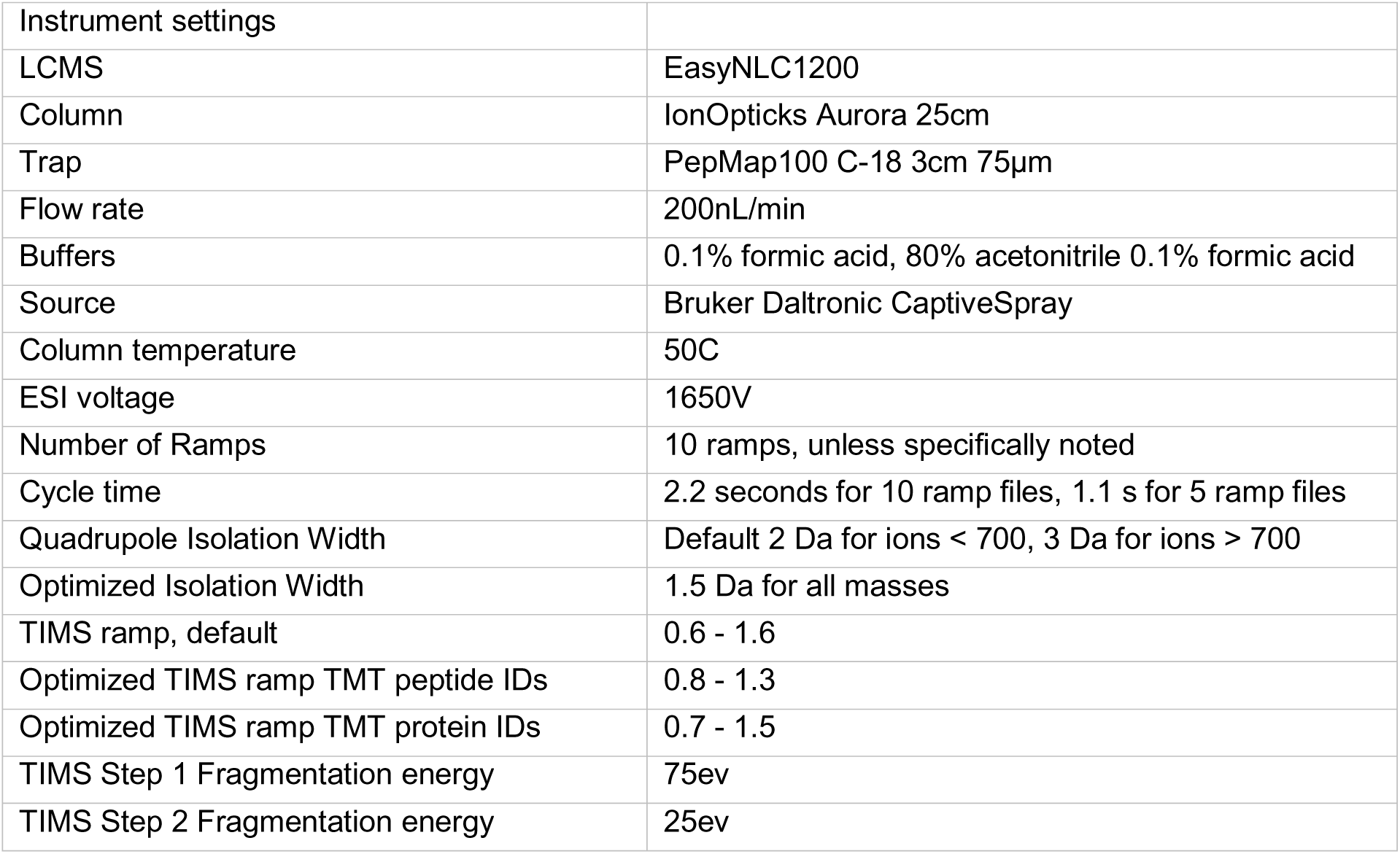
A summary of the instrument settings used in this study

### Data conversion and processing

Vendor proprietary output (. d) files were converted to MGF using three separate solutions: ProteoWizard MSConvert Developer Build Version: 3.0.20310-0d96039e2 for both GUI and command line-based conversions. MSConvert parameters for default pasef MGF conversion were utilized through the GUI. MSConvert combined MS/MS spectra if all of the following criteria were met elution time ≤ 5 seconds, ion mobility tolerance ≤0.1 1/k0 and precursor m/z ≤0.05 Da. Noise filtering was performed through the MSConvert via command line using the same parameters with additional filtering for the 200 most intense fragment ions from each MS/MS spectra. FragPipe 14.0 was used for the direct analysis with MSFragger 3.1, Philosopher 3.3.11 and Python 3.8.3.23 All calibrated MGF spectra were generated with MSFragger 3.1.1. which was released during the construction of this manuscript and enabled the recalibration functionality. All MSFragger settings were set at default for pasefDDA data closed search with the addition of the TMTPro reagents as dynamic modifications on the peptide N-terminus and on lysines. Conversion of data through Data Analysis 5.3 was performed via a Visual Basic script provided by the vendor.

### Processing of pasefRiQ data in Proteome Discoverer

All MGF files were processed with Proteome Discoverer 2.4 using SequestHT and Percolator with the reporter ion quantification node. A scan filter was used to the unrecognized fragmentation of the TIMSTOF MS/MS spectra as HCD and the spectra as FTICR at a resolution of 35,000 in order to allow the visualization of all significant figures in downstream processing. To reduce the complexity of TIMSTOF MS/MS spectra a binning method was used that retained only the 12 most intense fragment ions from each 100 Da mass window within each spectrum. Spectra were searched with SequestHT using a 30ppm MS1 tolerance and a 0.05 Da MS/MS fragment tolerance. Static modifications of the corresponding TMT reagent and the alkylation of cysteines with iodoacetamide was employed in all searches, with the exception of PTM searches in which the TMTPro reagent PTMs on lysine were added as dynamic modifications. The UniProt SwissProt reviewed FASTA was used for the corresponding organisms from downloads of the complete reviewed SwissProt library in January of 2021. The cRAP database (www.gpm.org) was used in all searches. For quantification of the TMTPro9 samples, a custom quantification scheme was built from the TMTPro defaults that disabled the C13 isotopes of each pair. The vendor default workflow for reporter-based quantification was used with the following alterations: only unique peptides were used for quantification, unique peptides were determined from the protein, not protein group identification, the default quantification scheme was intensity based, and the minimum average reporter intensity filter was set at 10. The protein marker node was used to flag and filter contaminants from the cRAP database and the result statistics node was added for post-processing. Carrier proteome data was analyzed both with and without vendor supplied impurity data. As no meaningful alterations were observed when these were applied at these high dilution levels, these were not applied or considered for biological data. Data was viewed using the IMP-MS2Go (www.pd-nodes.org) release for Proteome Discoverer 2.5.27. Single cell population subgroup analysis to study cells in individual cell cycle stages and for reanalysis of treated cells expressing measurable levels of CLIC3 was performed in PD 2.4 by creating new study factors and manually relabeling individual cells for quantitative analysis.

### Processing of pasefRiQ single cell data in MaxQuant and Proteome Discoverer

The sixty-three LCMS runs, containing a total of 441 single cells from this study, vendor proprietary. d files were converted to MGF using MSConvert using the default DDA pasef conversion method. The calibrated MGF files were processed in Proteome Discoverer 2.4/2.5 as described above. Percolator was used for FDR estimation, as well as for FDR estimation at the peptide and protein group levels. For normalization, the files were reprocessed with identical settings in two separate workflows. The first had the removal of the 134N and 133N reporter channels. This allowed the comparison between the signal and blank (126) channels but did not permit normalization due to the scaling of the signal in the blank channel. A second normalization and scaling using a total sum-based approach was used for final analysis. Entire LCMS runs or individual sample channels were removed from consideration as outliers in downstream data analysis when no reporter ion signal was obtained that exceeded that of the blank method blank channel. In addition, the 133N channel was excluded from all analysis due to significant ratio distortion which was identified as impurities in the 134N carrier channel from the TMT reagent kit. For MaxQuant analysis, the recommended settings described for pasefDDA 64 were used in version 1.6.17, with the addition of the 9 TMTPro tags which were manually added to the XML schema. For PTM analysis the spectra were searched with SequestHT in Proteome Discoverer 2.4 with lysine acetylation and the phosphorylation of serine and threonine considered as possible modifications. The IMP-PTMrs node was used for modification site localization scoring. For TIMSTOF data a precursor and fragment tolerance of 30ppm was used for all analyses.

### Reporter ion filtering and database reduction

We have recently described the development and implementation of an automated pre-analysis quality control software for multiplexed single cell proteomic analysis.^64^ DIDARSCPQC allows the end user to specify quality conditions for filtering MS2 spectra for downstream analysis. In addition, the program provides output metrics to flag cells within multiplexed sets which have failed analysis. We used DIDAR to filter the converted and calibrated MGF files from this study by requiring that at least one reporter ion corresponding to a well containing a single cell in wells 127n-131n. The MS2 spectra was moved to a new file with the prefix “Filtered” applied to the MGF file name if an ion was detected within a mass tolerance of 0.005 Da of the exact mass of the reporter ion. This round of filtering reduced the total number of MS2 spectra from approximately 2.2 million to 1.4 million. Following manual review of the DIDAR output we chose to remove entire files from consideration that did not exhibit more than 3x the number of spectra observed with a 126 method blank control signal using the same criteria. The remaining filtered LCMS runs were used for downstream analysis for PTM identification.

### Pathway and gene ontology analysis

Three commercial programs were utilized for pathway analysis, Protein Center (Thermo) and Ingenuity Pathways Analysis (Qiagen). For protein center analysis, differential proteins were selected from the normalized data in proteome discoverer and all proteins not meeting a cutoff of 2-fold at a p value <0.05 were excluded. The top pathways were determined by the number of remaining proteins that were identified within that group. For IPA analysis, the normalized ratios of all proteins with quantification of sotorasib/control were exported as CSV and uploaded into IPA using core analysis. The following settings were used, Core Expression Analysis based on Expr Fold change utilizing z-scores for directional analysis. Files were compared against the Ingenuity Knowledge Base at the gene level. Only experimentally observed relationships were used for pathway construction with filtering for human samples and cell lines. The SimpliFi cloud server (Protifi, Toronto, Canada) beta version was used for downsteam analysis and visualization of all data using the default interpretation settings for Proteomics data and a direct import of the MaxQuant output file. For CLIC3 positive cell analysis, output UniProt identifiers and relative fold changes were copied as .txt and uploaded into StringDB and matched against the default human database. All annotations meeting default cutoffs are provided as **S.Data 11**.

### Phosphopeptide quantification

Identified phosphopeptides were filtered using a log2 fold change of 1 using Boolean logical filters within Proteome Discoverer 2.4 and a minimum peptide confidence cutoff filter of approximately 0.05% and a localization confidence score from phosphoRS of 70%. This final list consisted of 1435 phosphopeptides, which were exported to CSV for normalization. The ratios of all proteins containing these phosphopeptides in addition to >1 unique unmodified peptide, as determined from the ungrouped protein level were exported to .csv. Phosphopeptide ratios were normalized against the total protein ratio of the 673 proteins they mapped to through use of an in house developed tool provided with this manuscript. Normalized phosphopeptides with ratios demonstrating a log2 fold change of greater than 2 were exported to CSV for pathway analysis.

### Comparison of scSeq and proteomics data

The mean of the normalized and transformed transcript abundance from the scSeq data of all H358 cells treated with sotorasib was used as proxy to simulate bulk RNASeq transcript abundance. The ratio of each mean transcript abundance was calculated and the transcripts ranked. For visualization of TOP2A transcript expression, the abundance for each individual cell was converted to a 3 dimensional matrix using a custom tool where a cell number and treatment condition composed the x and y dimensions and transcript abundance was plotted in the third dimension. The protein expression data was log scaled and converted to the same three dimensions and the data was combined and plotted in the open GlueViz environment. A visual basic converter capable of converting both SCP and scSeq data to this three dimensional format is made publicly available at https://github.com/orsburn/gluevizSingleCell. Two dimensional plots were assembled natively and the three dimensional viewer utilized through the VisPy plugin. All analysis was performed in Anaconda 1.9.12 and GlueViz 1.0.0.

## Code and Availability

A summary of the Proteome Discoverer results for proteins and phosphopeptides can be downloaded at this direct link. Phosphopeptides ratios were normalized against protein abundances using LPN V1.0 which is available at https://github.com/orsburn/LazyPhosphoNormalizer. All copy number distribution plots were generated using the openly available Shiny tool, available at https://proteomicsnews.shinyapps.io/CopyNumber_plotterV2/. Comparison of single cell proteomics and scSeq data was performed using the tools provided at https://github.com/orsburn/gluevizSingleCell. The custom database and XML aspects used for the analysis of cell cycle status of individual cells are available at: https://github.com/orsburn/SCP_cell_cycle_stripping. All additional figures were generated in R Studio 1.3.1093 utilizing Tidyverse, ggplot and ggally using the most up to date versions available on 3/10/2021.

## Data and Availability

Results from previous SCoPE-MS studies utilized in this study are available at www.ProteomeXchange.org via the following accession numbers: PXD008985, PXD011748 and PXD025387. Optimization files for background coisolation reduction and two proteome standards at 240ng are available at the following link: ftp://MSV000088757@massive.ucsd.edu with reviewer password: pasefRIQ. Following publication, the data will be permanently available at: ftp://massive.ucsd.edu/MSV00008875. Files used for optimization of pasefRiQ at dilution levels relevant to single cell loads are available at this link: ftp://MSV000088796@massive.ucsd.edu using the password: pasefRiQ. All files and processed data used for the calculation of the carrier proteome effect in this study are available at: ftp://massive.ucsd.edu/MSV000089428/. All vendors. d files, MGF peak lists used in the study, MaxQuant protein lists and Proteome Discoverer results for protein and phosphopeptide identifications in H358 single cells are available through ProteomeXchange as PXD028710. During review, these can be accessed via using reviewer credentials: MSV000088144 and Password: pasefSCOPE700N. Direct download is available through ftp://MSV000088144@massive.ucsd.edu. In addition, all filtered MGF files used for MSFragger open search, mutational assignment and processed data are available at MASSIVE through: ftp://MSV000088157@massive.ucsd.edu with reviewer credentials: MSV000088157 and password: pasefSCOPE700N. The Proteome Discoverer output demonstrating all proteins and PTMs identified by MSFragger, MSAmanda and Sequest is available at: https://doi.org/10.6084/m9.figshare.19749745.v1. Finally, MS2Go hyperlinked output sheets containing statistics of all peptides, proteins and decoy data from single cells as well as an HTML interface to directly access all peptide spectral match data for all PTMs reported in this work are available at this link: https://doi.org/10.6084/m9.figshare.19749751

## Acknowledgments

We would like to thank Megan Rigby for advice on the use of sotorasib in cell culture and Dr. Alexis Norris for assistance with custom R tools featured in this work. In addition, we would like to thank Dr. Amol Prakash for access to, and assistance with, the Bolt cloud search engine.

## Inventory of supplemental data

**S.Data 1. Data related to Fig. 1A/B** and **Fig. 2A/2B**. A summary of optimization experiments to develop pasefRiQ using the TMTTKO commercial digest standard and an in-house developed 2- proteome standard digest model.

**S.Data 2. Data related to Table 1, Figure 2** and **S. Fig 10**. A comparison of a pilot analysis of H358 single cells compared to publicly available files from the SCOPE2 study (Sheet1). The dilution series used to test the carrier proteome effect in pasefRiQ (Sheet 2). Summary results of the TNAa protein at each carrier level.

**S.Data 3. Data related to Figure 3.** MS2Go output of all data obtained from single cells in this study using the MSAmanda, MSFragger and Sequest search engines with Percolator quality filtering.

**S.Data 4. Data related to S. Fig 11.** Proteins identified in single cells using 4 separate search engines. Sheet 1 is a composite of results from MSFragger, SequestHT, and MSAmanda 2.0 compiled with Percolator within the Proteome Discoverer 2.4 environment. Sheet 2 is the protein output text report from MaxQuant 1.6.17. The remaining sheets are single LCMS file comparisons using the first and last files of batch 1.

**S.Data 5. Data related to Figure 3C** and **S. Fig. 16**. A summary table of 119 cell cycle proteins and their relative levels of detection and abundance across single cells in this study. Related to Figure 3C.

**S.Data 6. Data related to Table 2.** MS2Go output sheet detailing all proteins, peptides and PSMs where PTMs were detected with MSFragger, MSAmanda and SequestHT using Percolator quality filtering and ptmRS for PTM localization.

**S.Data 7.** Summary output sheet from filtered MGFs processed through the O-PAIR search tool within the MetaMorpheus informatics environment containing all PSMs, decoys, and search settings used to identify glycopeptides and glycoproteins.

**S. Data 8. Data related to Fig 4C** and **4D**. A summary output sheet of proteins with PTMs detailing the localization site and total protein coverage.

**S. Data 9. Data related to S. Fig. 13.** Correlation coefficients of phosphopeptide abundance compared to proteins as expressed in single cells in this study.

**S. Data 10.** Panther pathway data in support of **Fig. 7C.** Panther pathway analysis of transcripts from single cell seq analysis of 1,000 control vs 1,000 sotorasib treated cells (Sheet 1) with a 2- fold differential cutoff. The same pathway analysis of 115 control vs 115 treated single cells by SCP (Sheet 2).

## Supplemental Figures

**S. Fig. 1.**
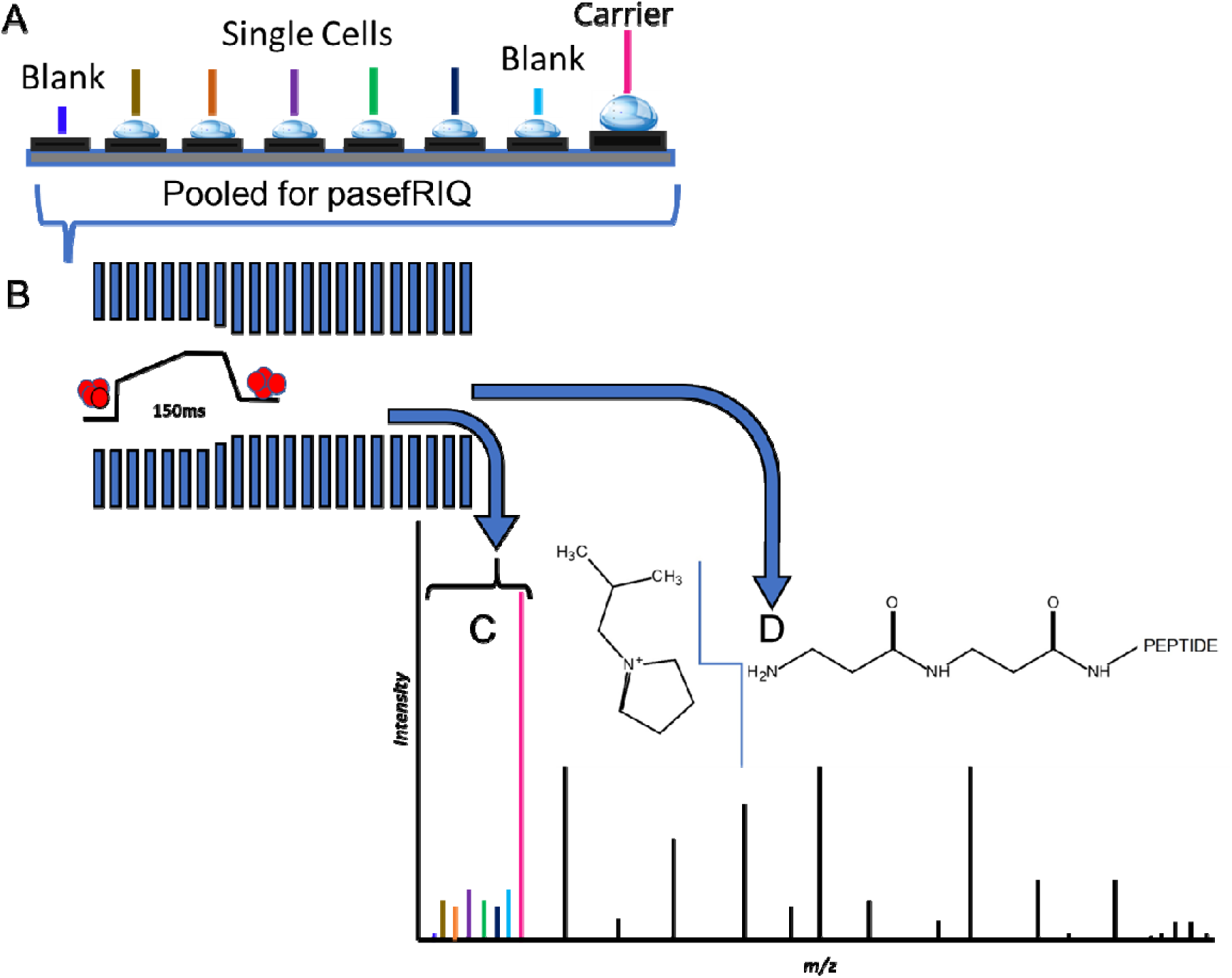
A cartoon illustrating the general strategy of analyzing single cells with pasefRiQ. A. Single cells and method blank control containing sorting buffer, but not a cell are sorted into wells of a receiving plate. Digested and labeled peptides are pooled for analysis. B. LCMS with a two stage pasef ramp is performed with ion accumulation occurring over a ramp time of 150 milliseconds. C. The resulting peptide is fragmented once using a high collision energy and pre- pulse trapping parameters optimized to capture low mass fragment ions for quantification. D. A second fragmentation even with lower collision energy and conditions to capture higher m/z fragment ions provide peptide sequencing information. The spectra from the two scans are combined in real time by the mass analyzer hardware.

**S. Fig. 2.**
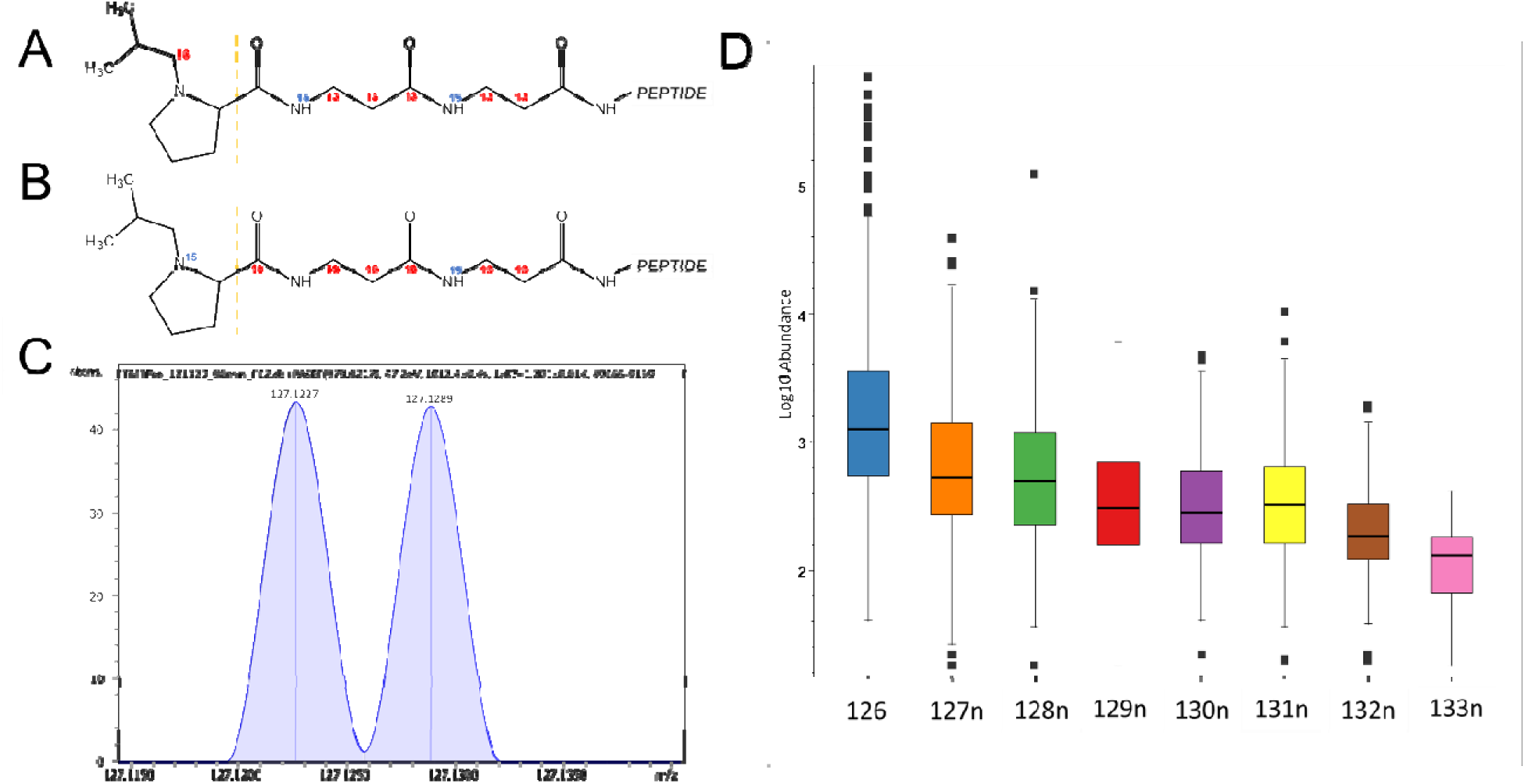
**A.** The structure of the 127N reporter ion from the TMTPro reagent. **B.** The structure of the 127C reporter ion. **C.** A representative image from the liberated reporter ion regions of a labeled peptide digest demonstrating that near-baseline separation of these ions can be achieved under some conditions. **D**. A boxplot representation of the dilution intensities presented in **Fig. 1B**.

**S. Fig. 3.**
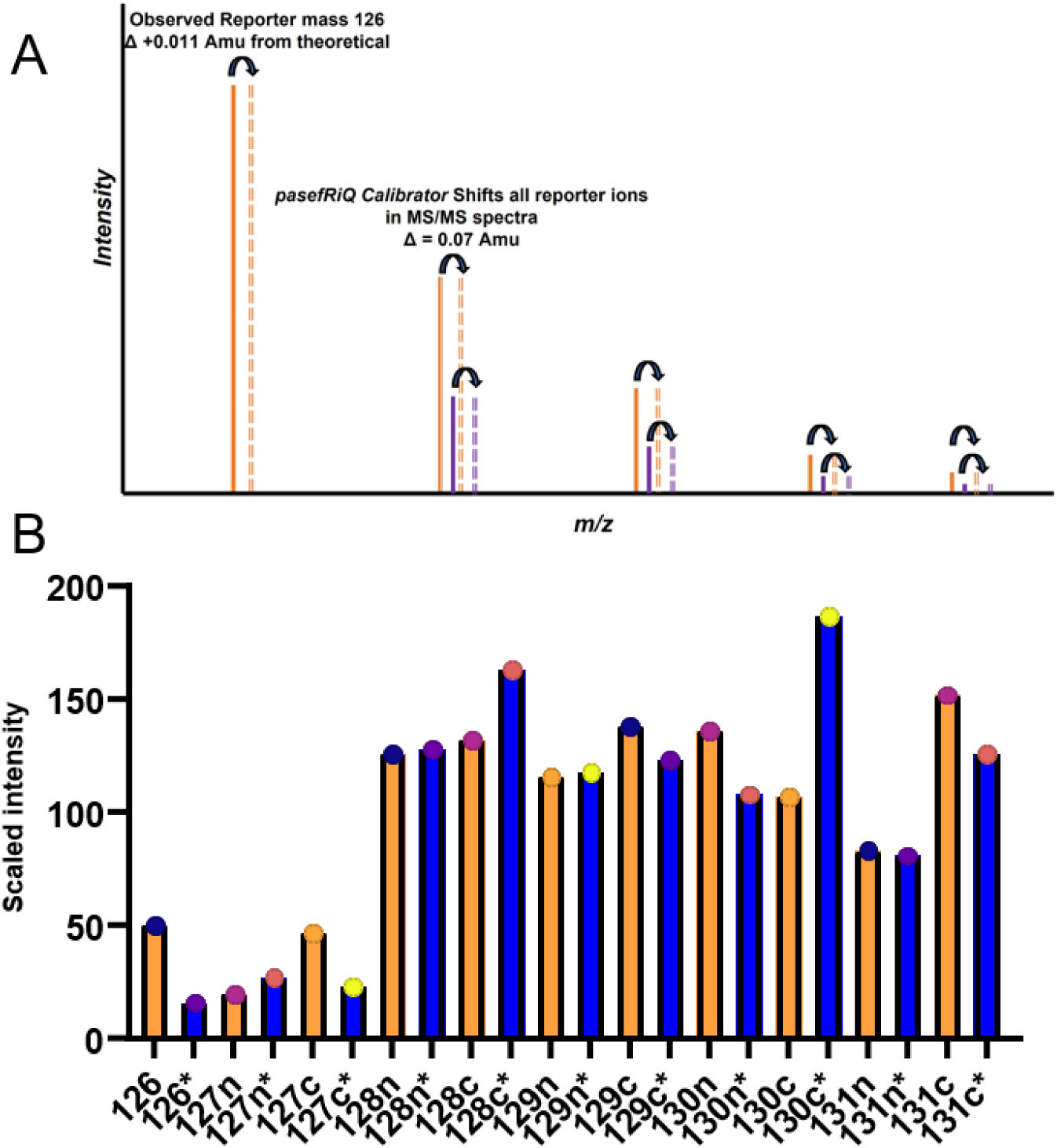
**A.** An illustration describing the MS/MS recalibration of reporter ions with the pasefRiQ Calibrator. **B.** The scaled intensity of the met6 protein in the TMT-TKO standard where * denotes the values following one point recalibration

**S. Fig. 4.**
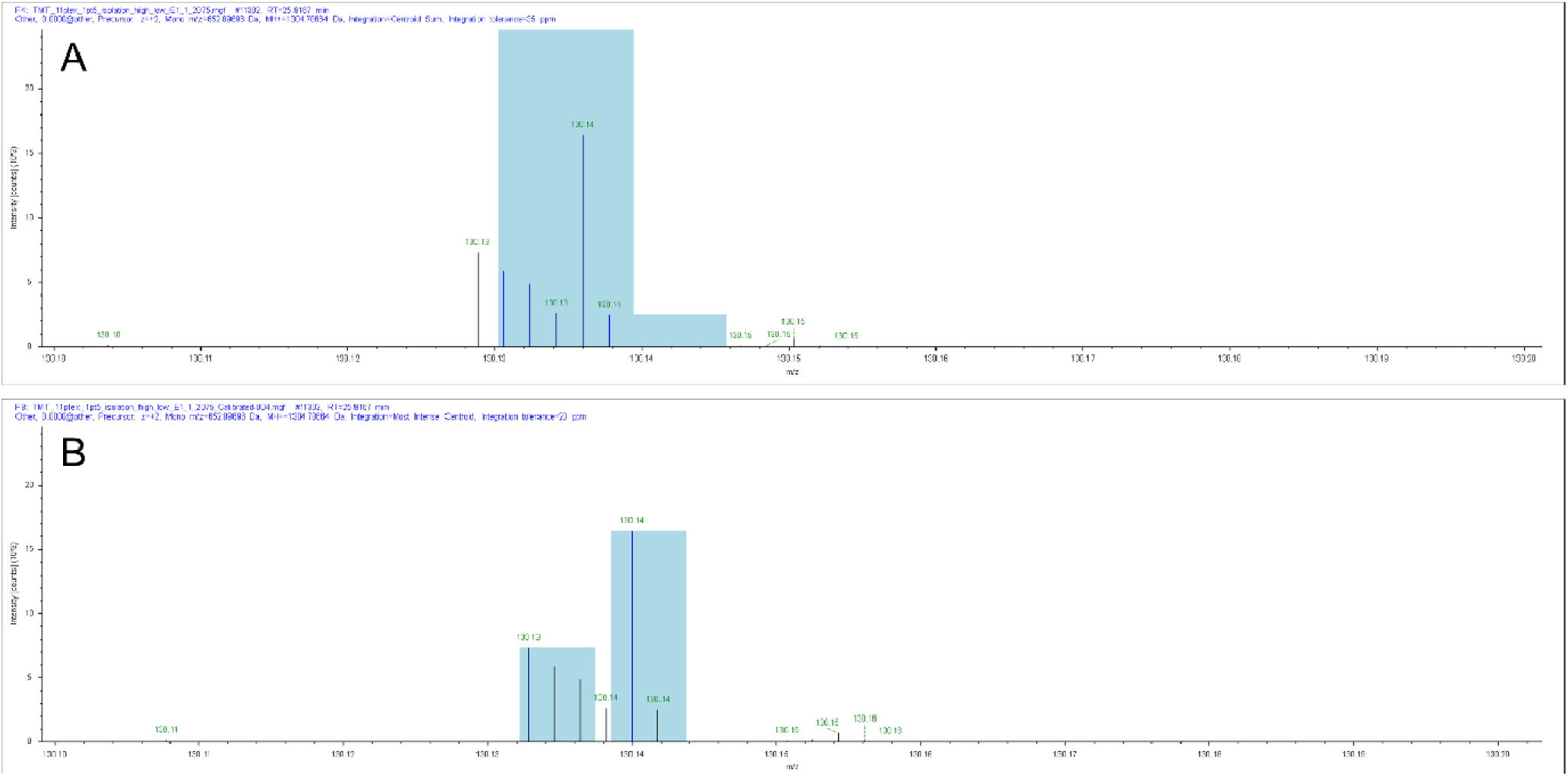
An example of the reporter ions integrated for quantification prior to and post recalibration **A.** VATSGVANK peptide reporter ion 130n and 130c with overlapping 35ppm integration windows displayed in blue. **B.** The same peptide following recalibration using a 20ppm non-overlapping integration windows.

**S. Fig. 5.**
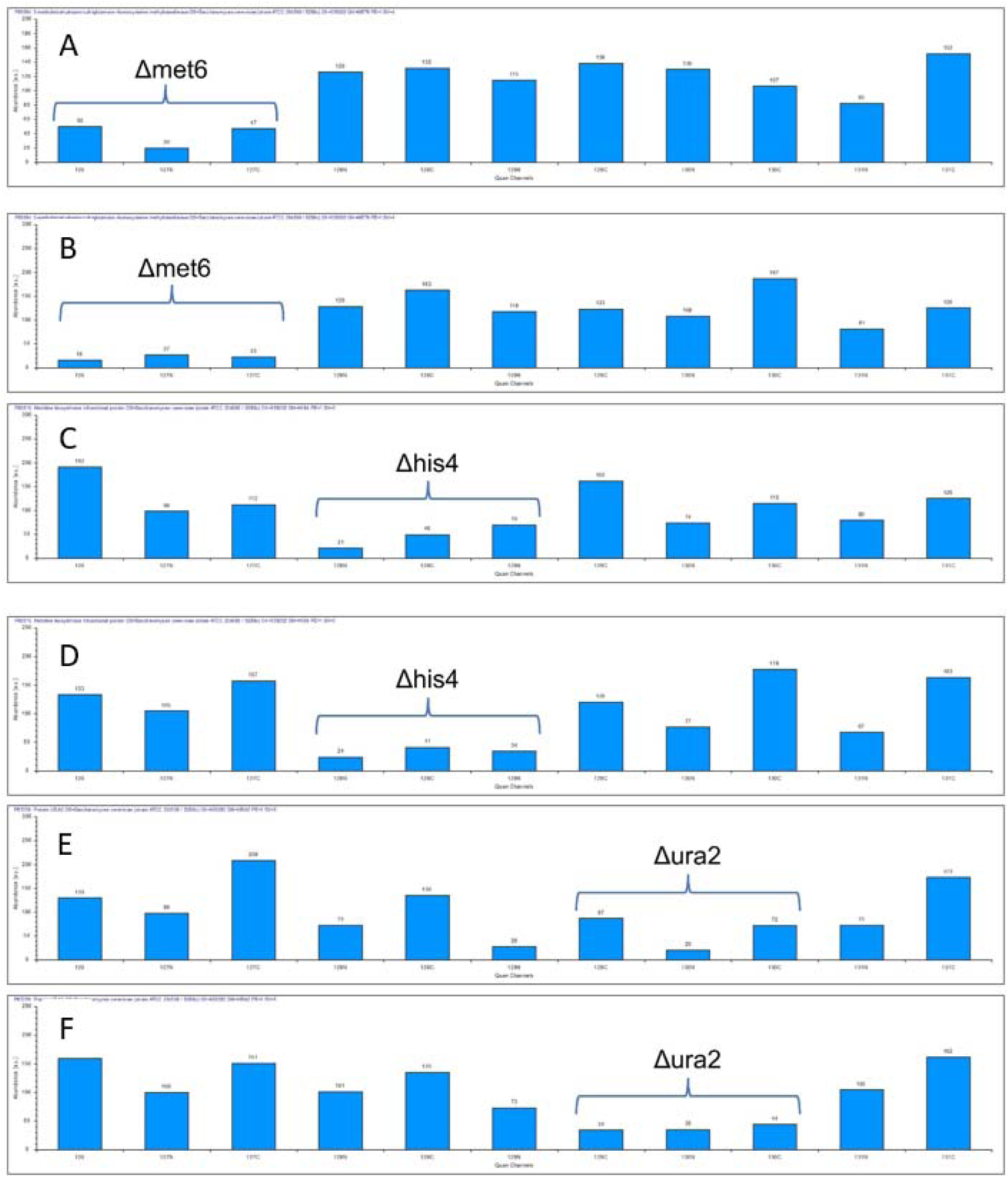
**A.,C.,E.** The scaled intensities of the Met6, His4, and Ura2 proteins from the TMT-TKO standard using an optimized pasefRiQ workflow integrated with a 20ppm mass tolerance window. **B.,D.,F.** The scaled intensities of the same respective files following a 0.004 Da recalibration with the *pasefRiQCalibrator*.

**S. Fig. 6.**
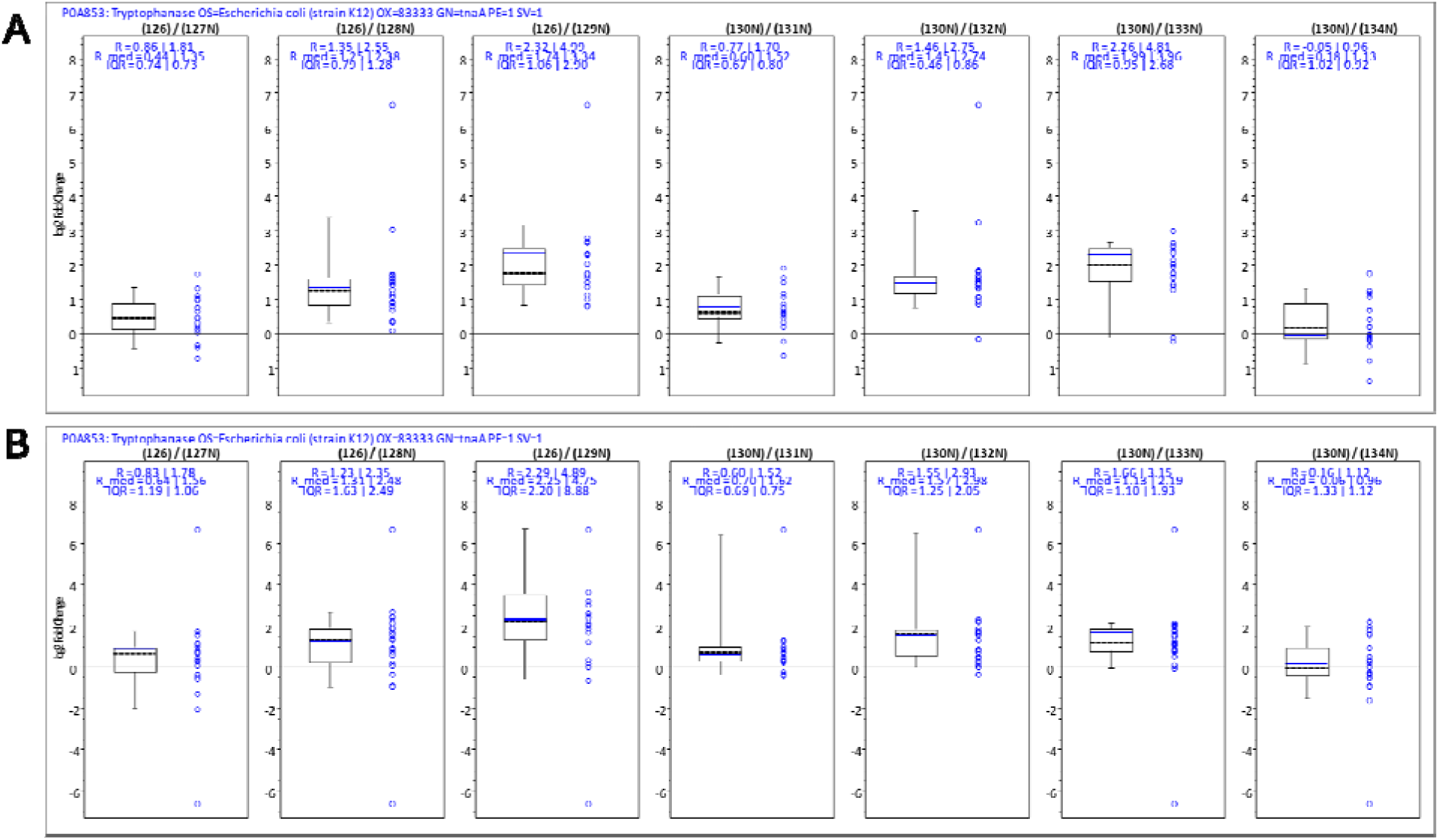
A. Boxplots demonstrating the raw peptide intensities for two of carrier proteome samples summarized in Figure 2D. A. The peptide intensity values for the lowest relative carrier load of 30x more than any other sample. B. The same for the highest relative carrier load approximating 180 times that of any other sample.

**S. Fig 7.**
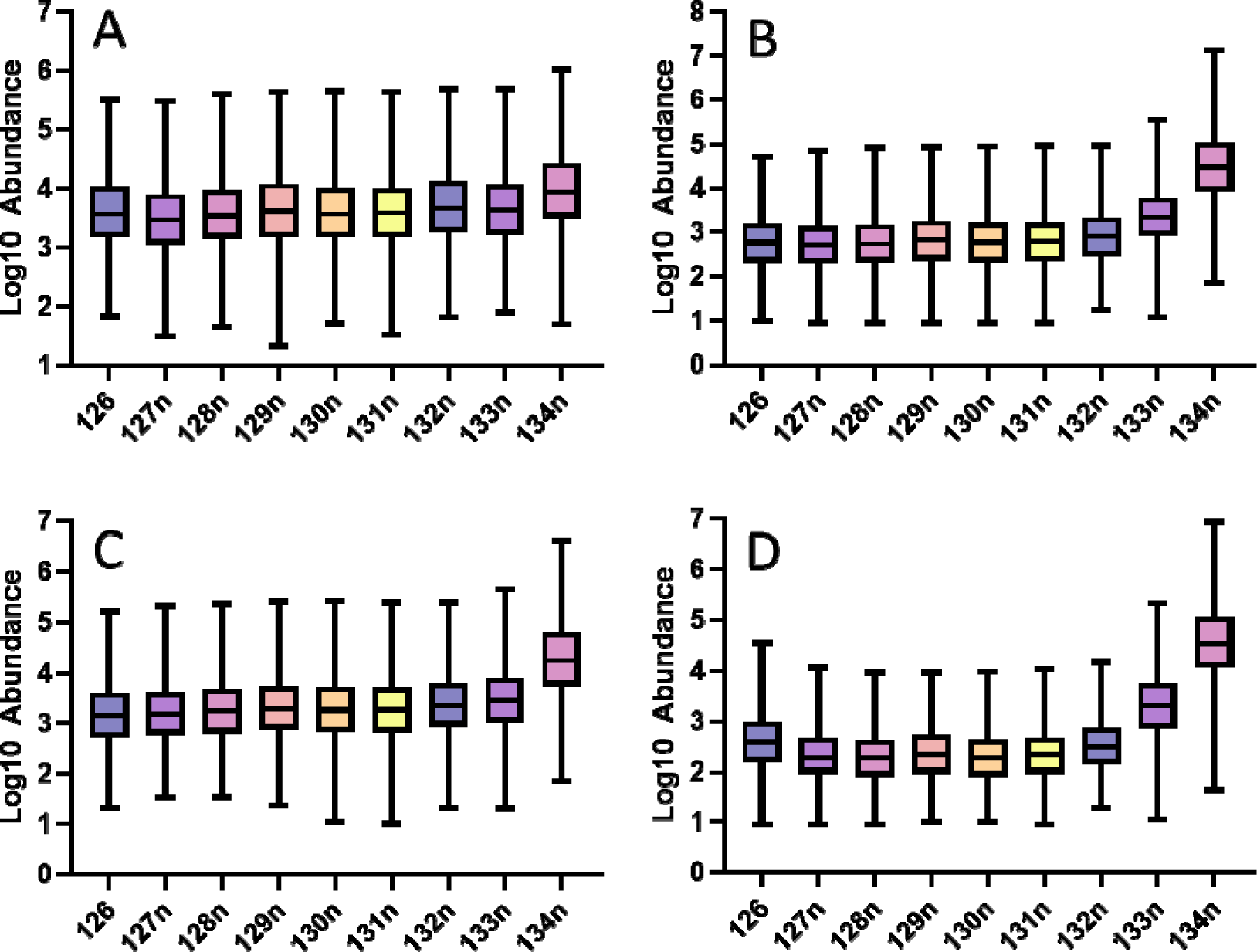
Box plots demonstrating the increasing effects of ratio distortion observed with increasing carrier channel loads: A. 1x carrier B. 50x carrier. C. 100x carrier. D. 500x carrier

**S. Fig 8.**
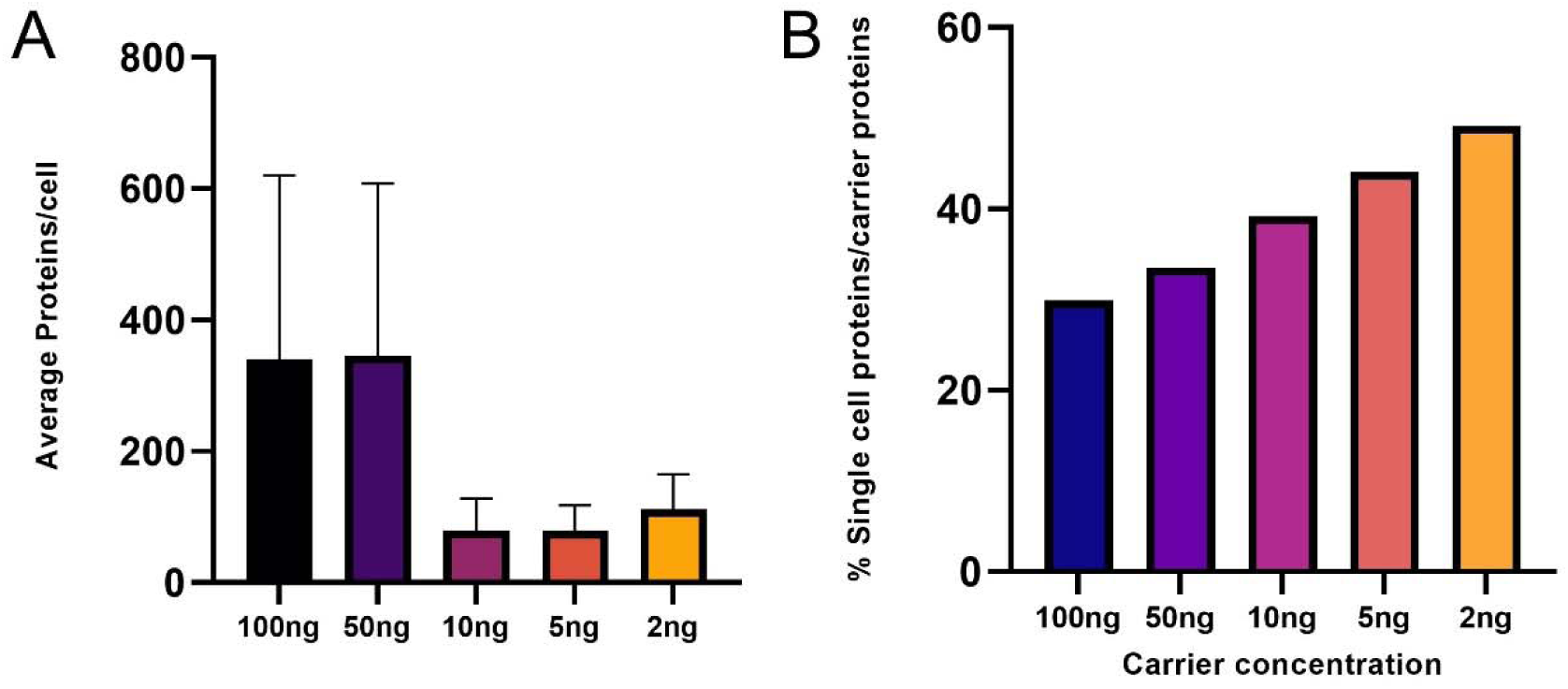
Bar charts representing the rough dilution series used to identify the ideal carrier channel concentration for single cells. **A.** Carrier loads of 100ng and 500ng provided comparable levels of verage protein identifications per cell. **B**. Lower concentrations of carrier corresponded to increasing signal from each protein per cell when compared to the identifications from the carrier channel.

**S. Fig 9.**
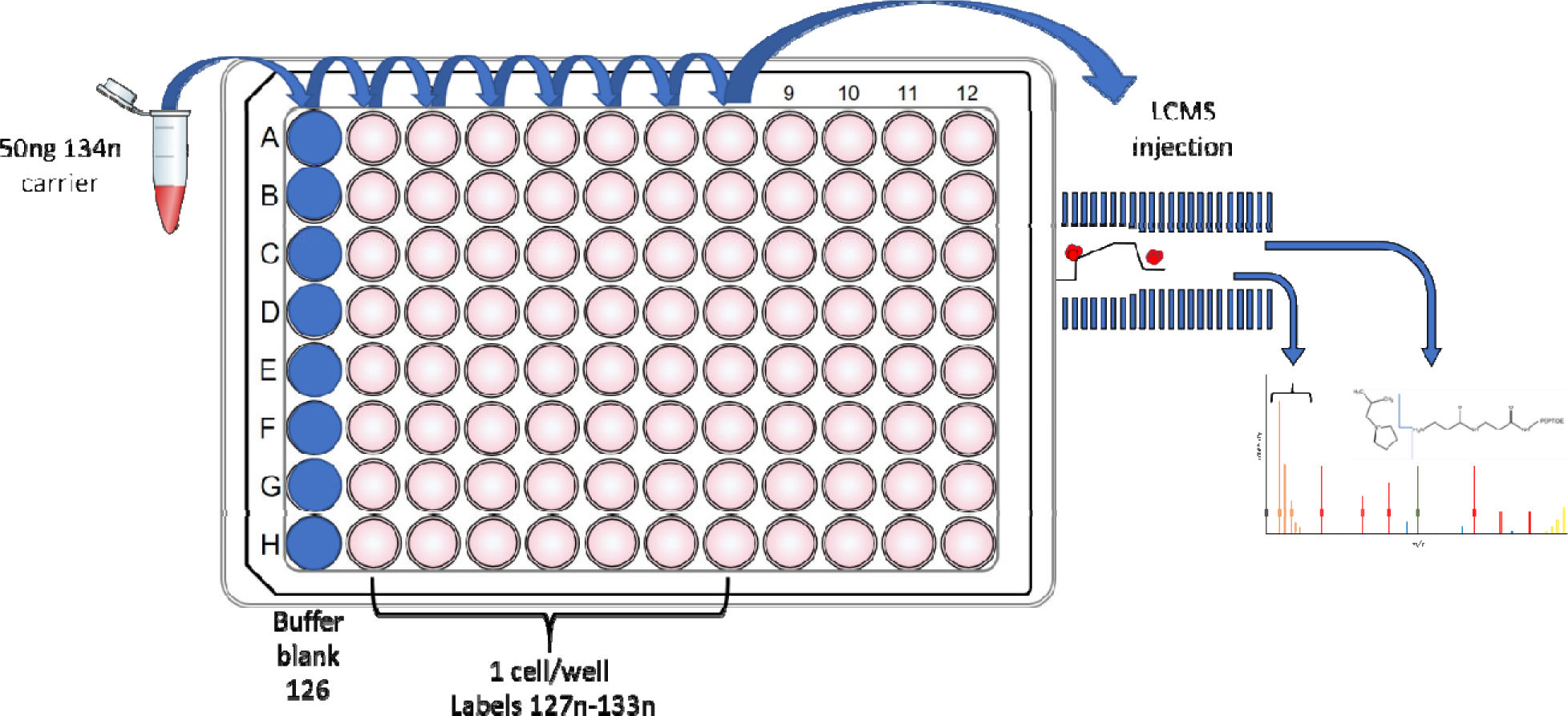
An illustration of the sample workflow for pasefRiQ analysis of single H358 cancer cells

**S. Fig 10.**
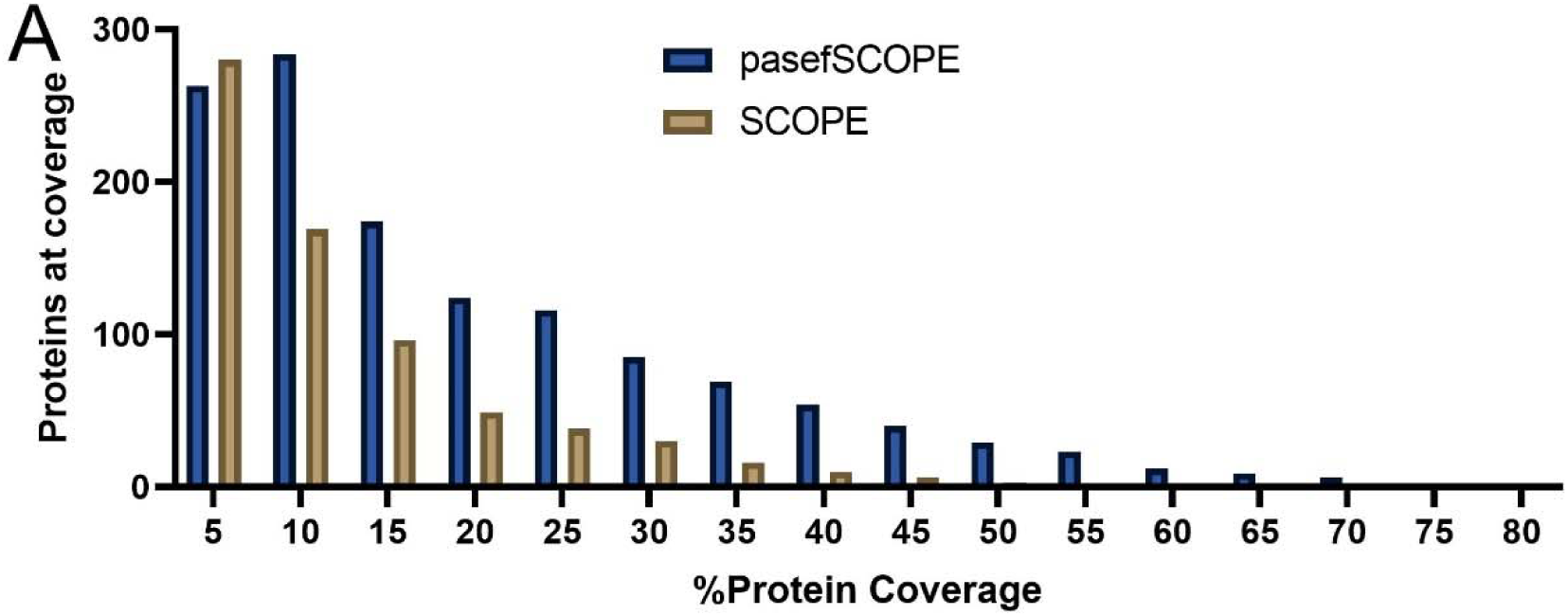
A plot comparing the relative percent sequence coverage of each protein identified in a pasefRiQ experiment compared to published SCOPE2 data using a D30 Orbitrap system.

**S. Fig 11.**
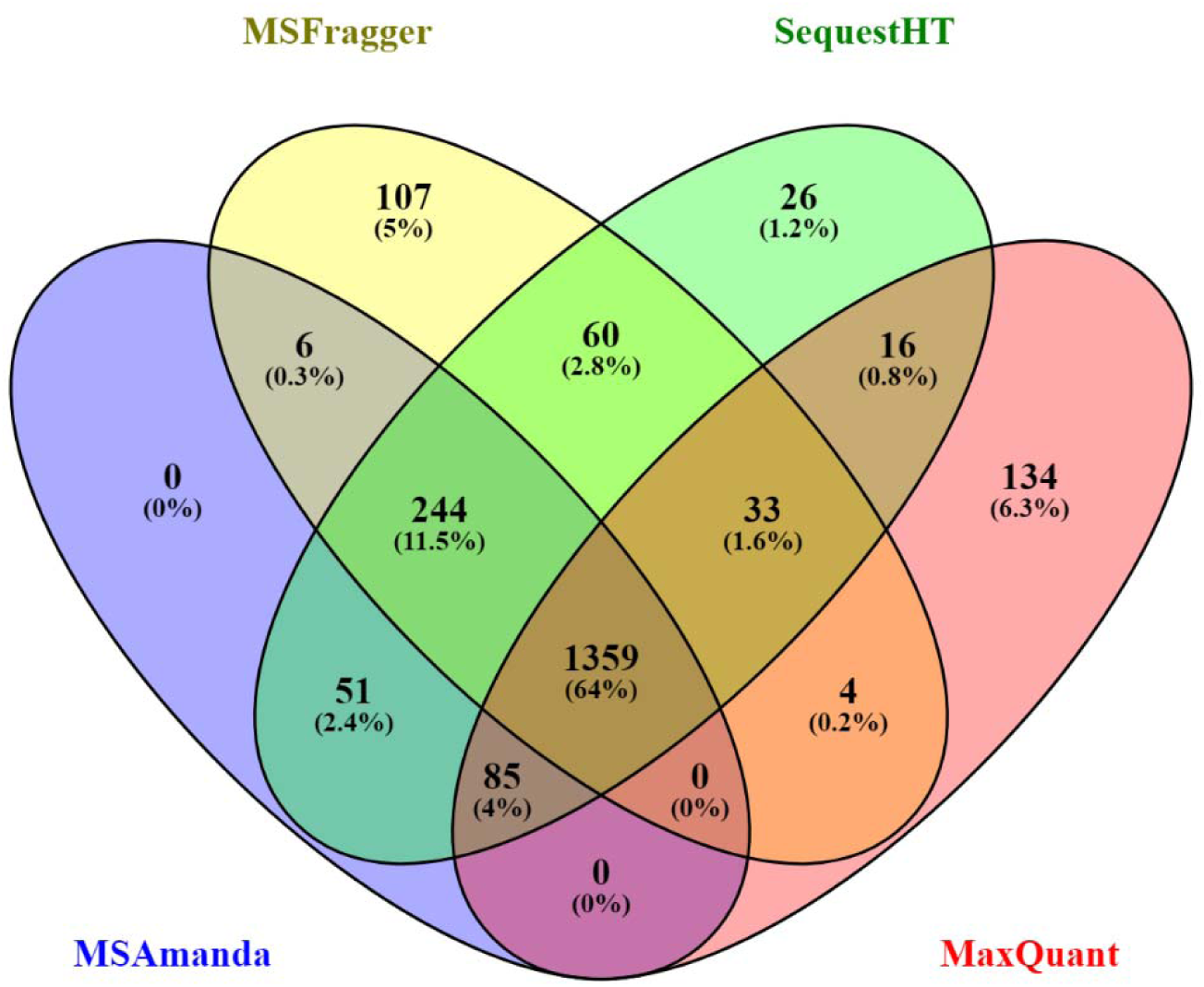
A comparison of the protein level identifications obtained by using four well- characterized search engines to process pasefRiQ H358 single cell data.

**S. Fig 12.**
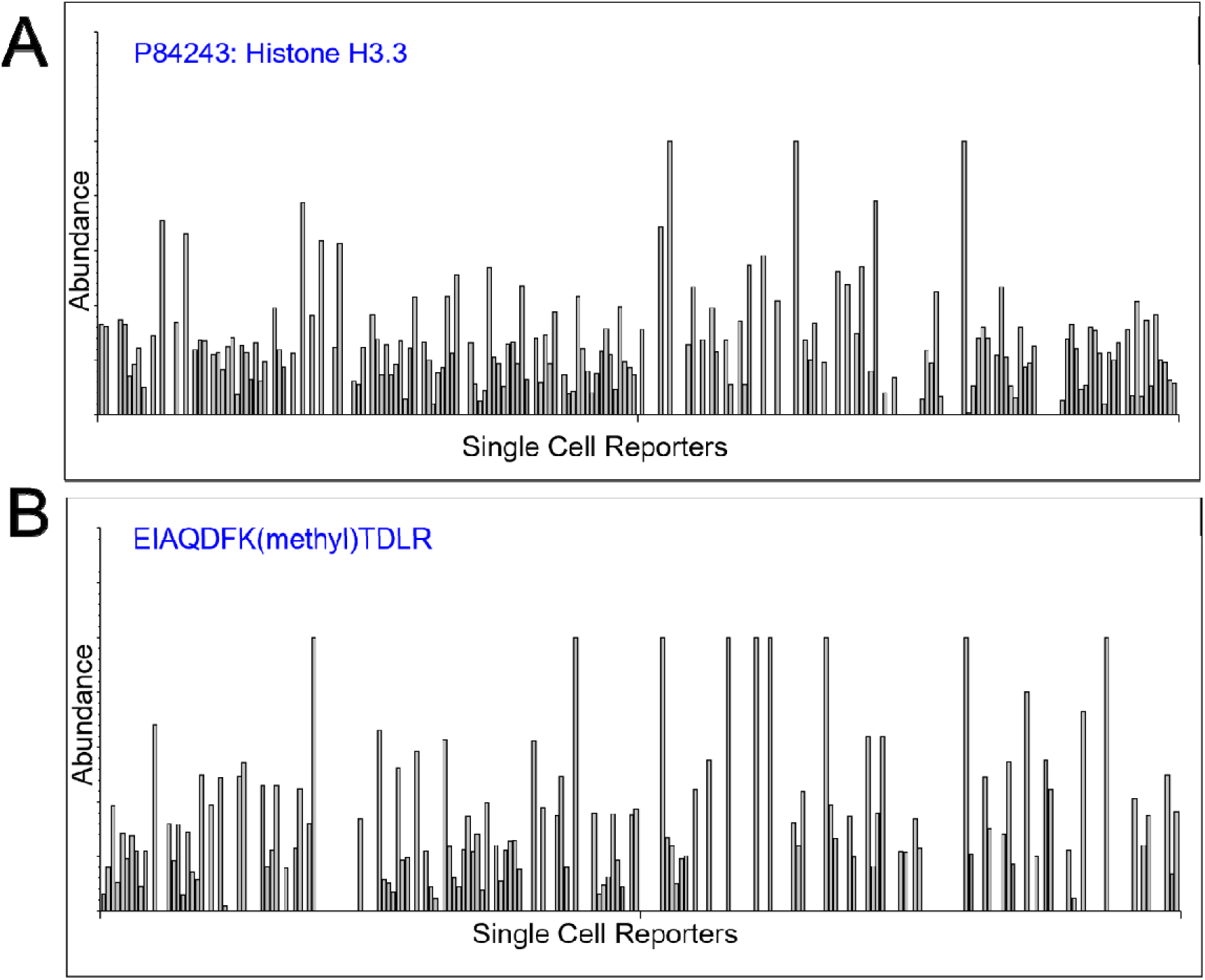
The relative abundance of histone proteins and PTMs across all single cells analyzed in this study. **A.** A bar plot demonstrating the identification rate and relative abundance of Histone 3.3 across 230 single H358 cells. **B.** A bar plot demonstrate the relative intensity of an identified methylation site on Histone 3.3.

**S. Fig 13.**
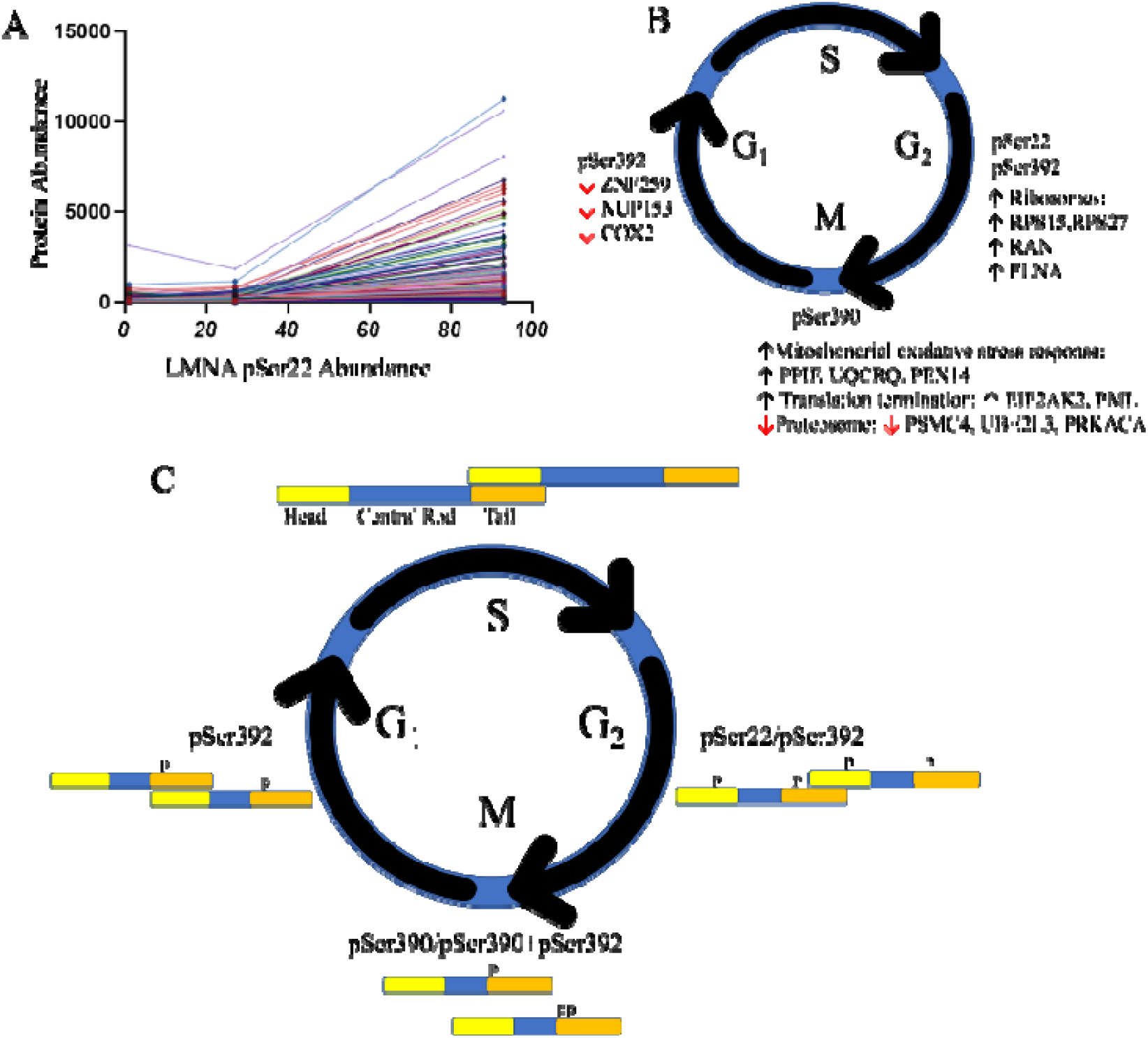
Correlation of LMNA phosphopeptides with other mitotic markers. **A.** Example correlation plots between LMNA Ser22 phosphopeptide abundance and proteins identified as cell cycle dependent by gene ontology analysis. **B.** A summary of peptides correlating with three LMNA phosphopeptides. **C**. A proposed mechanism of the activity of each phosphopeptide through the cell cycle of H358 cells.

**S. Fig. 14.**
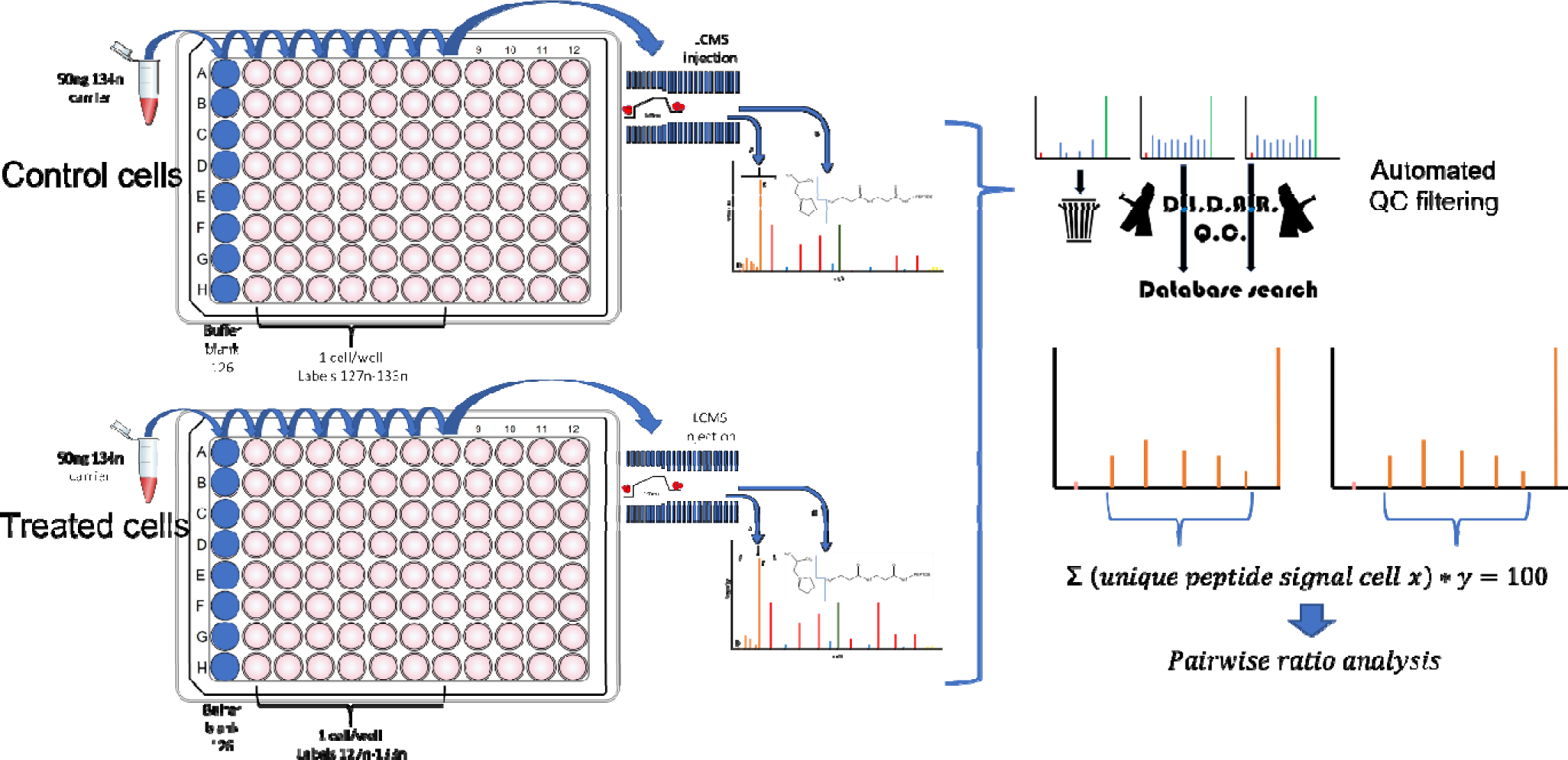
A cartoon demonstrating the overall experimental design, quality control and data analysis pipeline for the comparative analysis of 276 single cells in a drug treatment analysis.

**S. Fig. 15.**
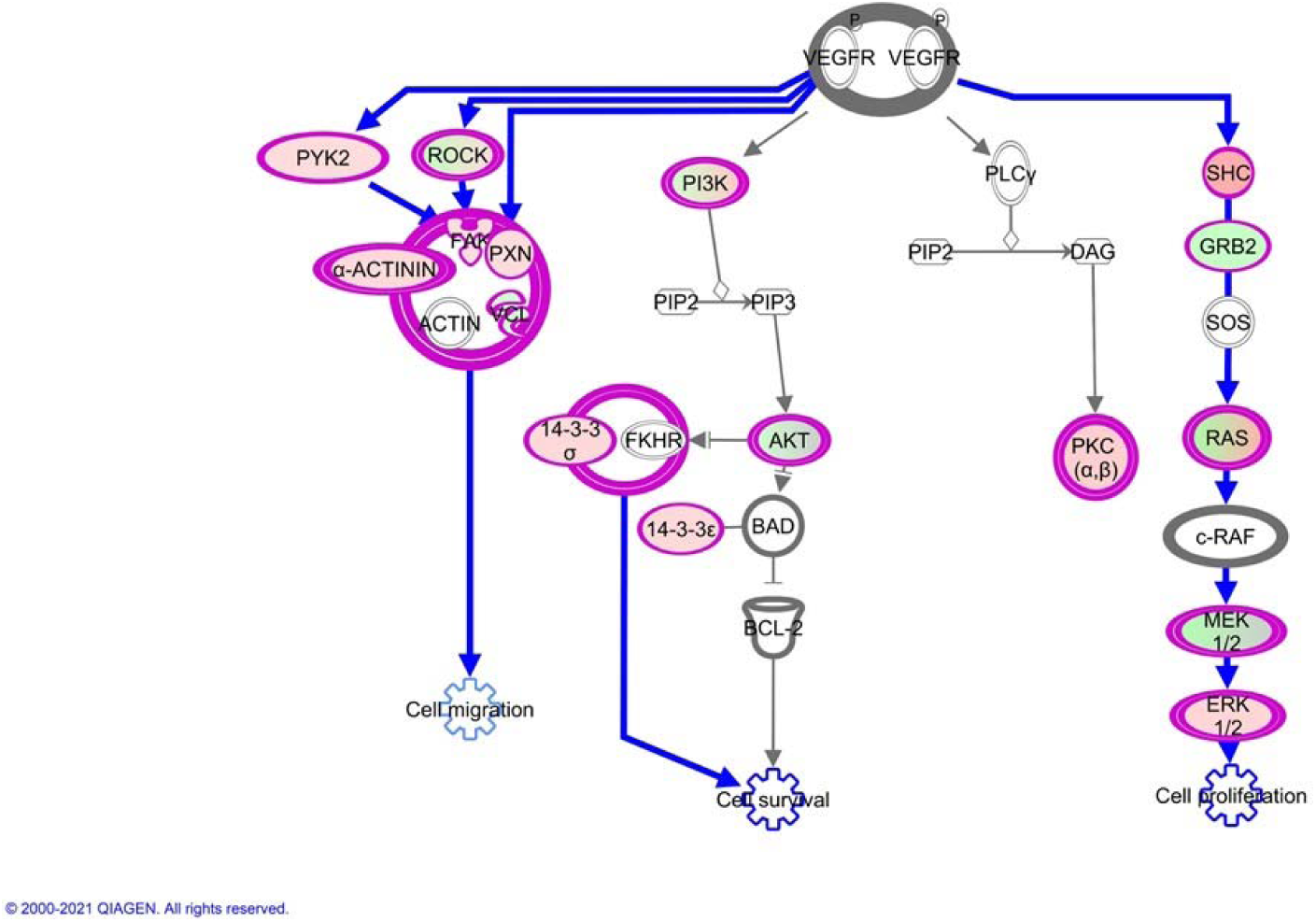
Pathway analysis indicated the VEGF pathway is the single most altered pathway in H358 cells following 40 hours of sotorasib treatment in single cells.

**S. Fig. 16.**
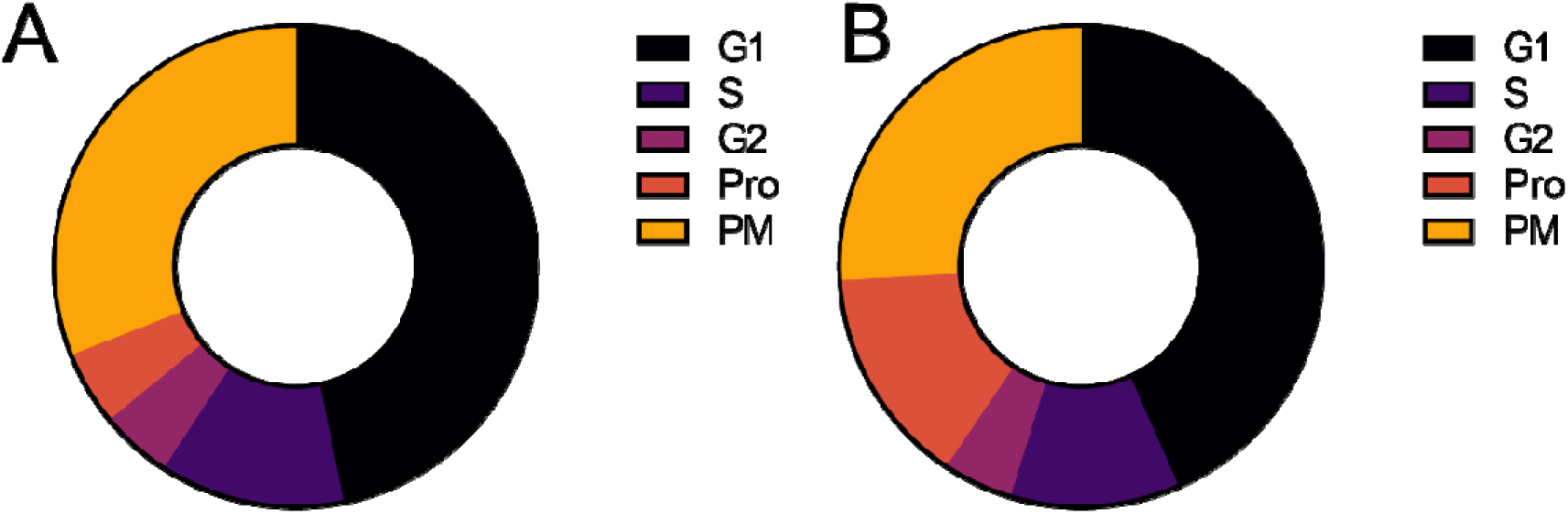
**A.** Shifts in cell cycle related protein abundance. **A** control H358 cells. **B.** Sotorasib treated cells

**S. Fig. 17.**
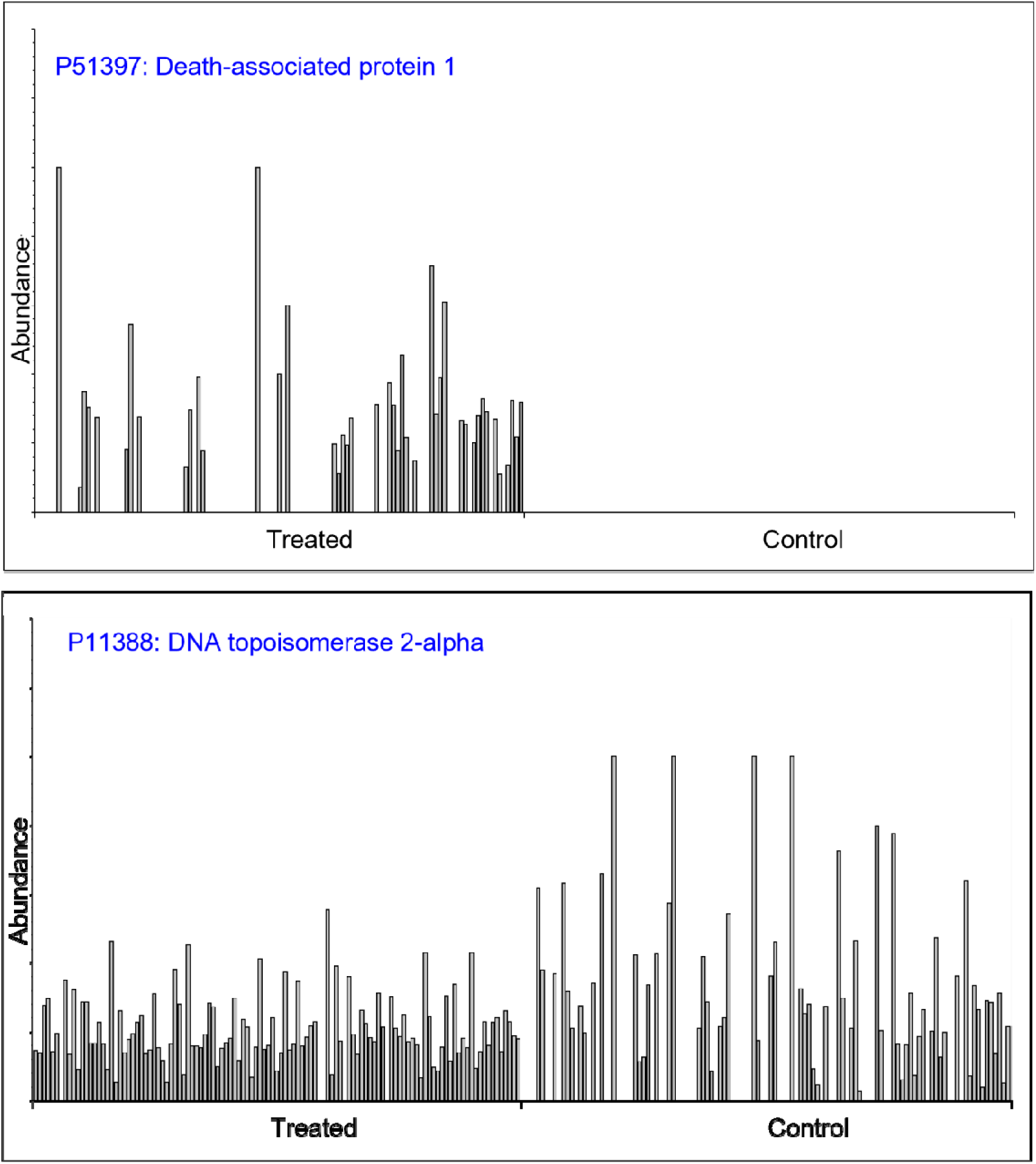
**A.** Visualization of the relative abundance of the DAP1 protein across control and treated cells demonstrating an observation that may be driven by a relatively small cellular subpopulation. **B**. The relative abundance of the TOP2A protein across cells demonstrating a mechanism that appears more relatively homogenous following drug treatment.

**S. Fig. 18.**
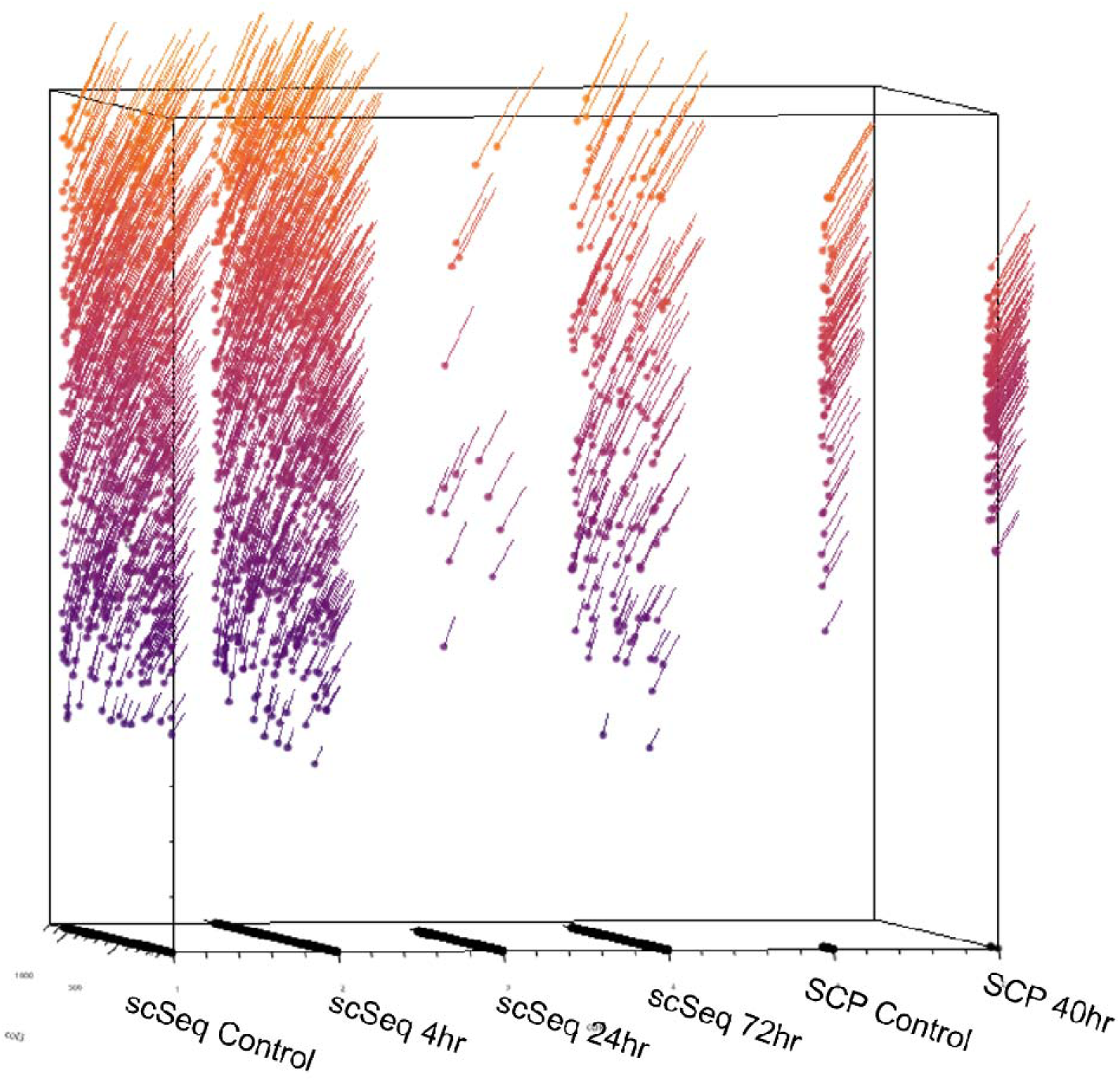
A comparison of the transcript and protein expression levels of TOP2A across approximately 4,000 and 230 single cells respectively.

**S. Fig. 19.**
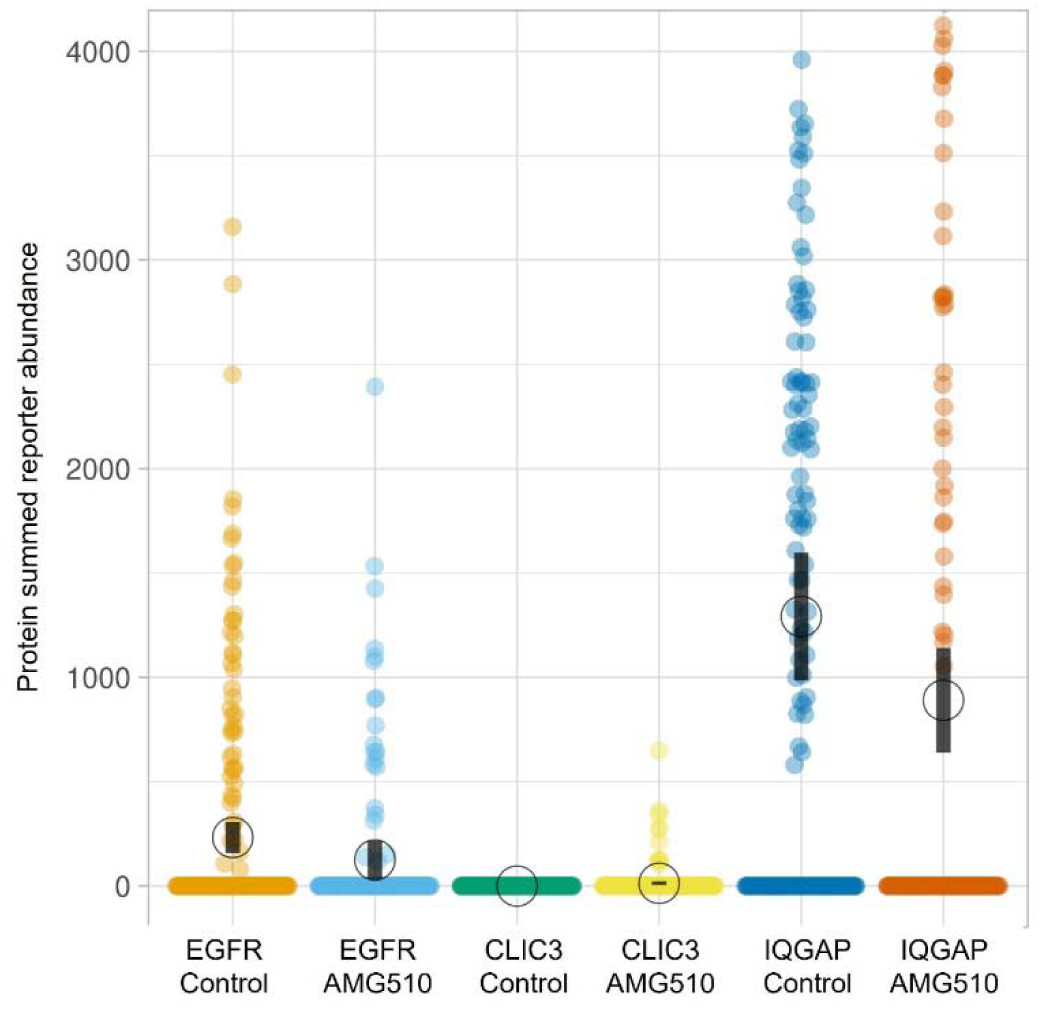
A visualization of the abundances of respective proteins across all single cells analyzed.

**S. Table 1.**
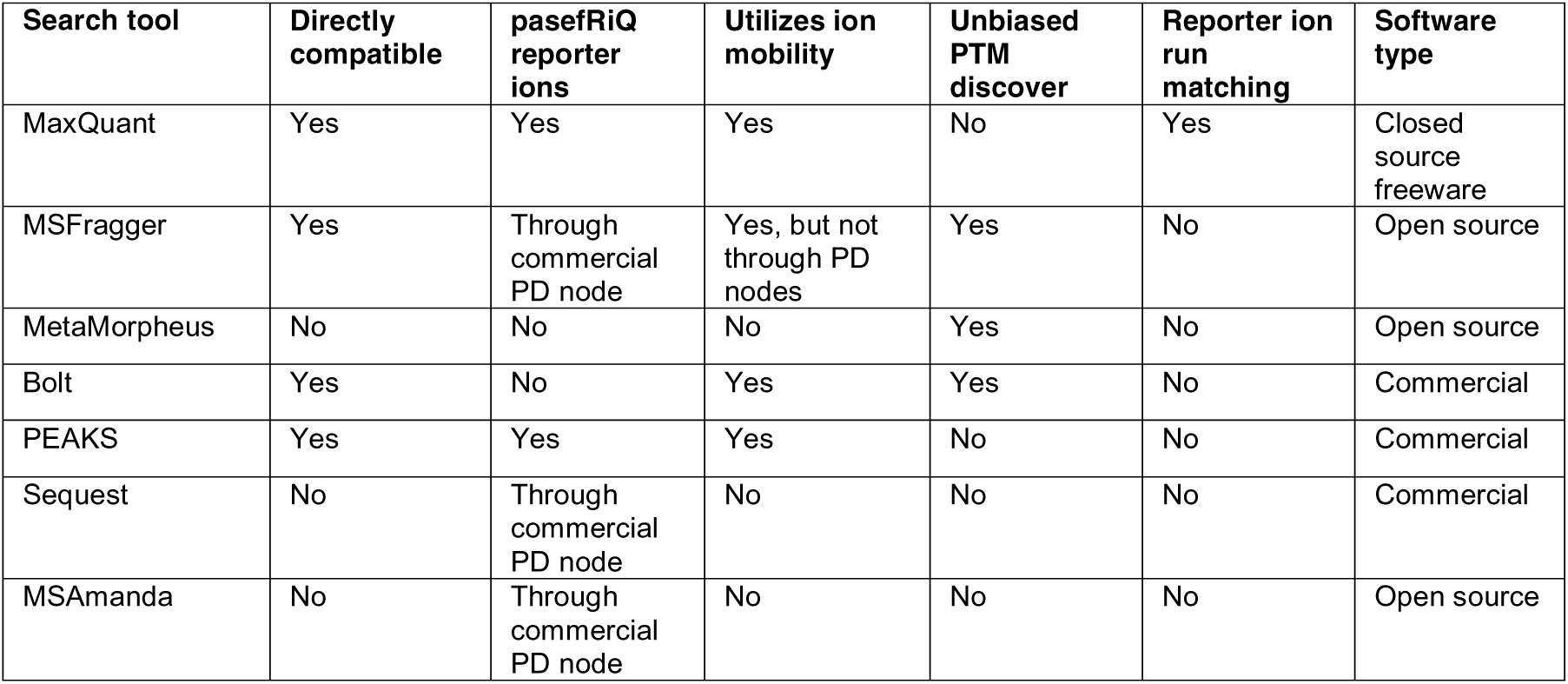
A summary of proteomics tools and their relative compatibility with TIMSTOF data files.

## References

1. Hwang B, Lee JH, Bang D. Single-cell RNA sequencing technologies and bioinformaticspipelines. Exp Mol Med. 2018;50(8):1–14. doi:10.1038/s12276-018-0071-8

2. Nagaraj N, Wisniewski JR, Geiger T, et al. Deep proteome and transcriptome mapping of a human cancer cell line. Mol Syst Biol. Published online 2011. doi:10.1038/msb.2011.81

3. Aebersold R, Agar JN, Amster IJ, et al. How many human proteoforms are there? Nat Chem Biol. Published online 2018. doi:10.1038/nchembio.2576

4. Slavov N. Unpicking the proteome in single cells. Science (80-). Published online 2020. doi:10.1126/science.aaz6695

5. Ctortecka C, Mechtler K. The rise of single-cell proteomics. Anal Sci Adv. Published online 2021. doi:10.1002/ansa.202000152

6. Hartlmayr D, Ctortecka C, Seth A, Mendjan S, Tourniaire G, Mechtler K. An automated workflow for label-free and multiplexed single cell proteomics sample preparation at unprecedented sensitivity. bioRxiv. Published online January 1, 2021:2021.04.14.439828. doi:10.1101/2021.04.14.439828

7. Specht H, Emmott E, Petelski AA, et al. Single-cell proteomic and transcriptomic analysis of macrophage heterogeneity using SCoPE2. Genome Biol. 2021;22(1):50. doi:10.1186/s13059-021-02267-5

8. Krug K, Jaehnig EJ, Satpathy S, et al. Proteogenomic Landscape of Breast Cancer Tumorigenesis and Targeted Therapy. Cell. 2020;183(5):1436–1456.e31. doi:https://doi.org/10.1016/j.cell.2020.10.036

9. Burke MC, Mirokhin YA, Tchekhovskoi D V., et al. The Hybrid Search: A Mass Spectral Library Search Method for Discovery of Modifications in Proteomics. J Proteome Res. Published online 2017. doi:10.1021/acs.jproteome.6b00988

10. Furtwängler B, Üresin N, Motamedchaboki K, et al. Real-Time Search-Assisted Acquisition on a Tribrid Mass Spectrometer Improves Coverage in Multiplexed Single-Cell Proteomics. Mol Cell Proteomics. 2022;21(4). doi:10.1016/j.mcpro.2022.100219

11. Tsai C-F, Zhao R, Williams SM, et al. An Improved Boosting to Amplify Signal with Isobaric Labeling (iBASIL) Strategy for Precise Quantitative Single-cell Proteomics *. Mol Cell Proteomics. 2020;19(5):828–838. doi:10.1074/mcp.RA119.001857

12. Hecht ES, Scigelova M, Eliuk S, Makarov A. Fundamentals and Advances of Orbitrap Mass Spectrometry. In: Encyclopedia of Analytical Chemistry. ; 2019. doi:10.1002/9780470027318.a9309.pub2

13. Meier F, Brunner AD, Koch S, et al. Online parallel accumulation–serial fragmentation (PASEF) with a novel trapped ion mobility mass spectrometer. Mol Cell Proteomics. Published online 2018. doi:10.1074/mcp.TIR118.000900

14. Gillson J, Ramaswamy Y, Singh G, et al. Small molecule KRAS inhibitors: The future for targeted pancreatic cancer therapy? Cancers (Basel). Published online 2020. doi:10.3390/cancers12051341

15. Zubarev RA. The challenge of the proteome dynamic range and its implications for in- depth proteomics. Proteomics. Published online 2013. doi:10.1002/pmic.201200451

16. Wiśniewski JR, Hein MY, Cox J, Mann M. A “proteomic ruler” for protein copy number and concentration estimation without spike-in standards. Mol Cell Proteomics. Published online 2014. doi:10.1074/mcp.M113.037309

17. Kaufmann A, Walker S. Comparison of linear intrascan and interscan dynamic ranges of Orbitrap and ion-mobility time-of-flight mass spectrometers. Rapid Commun Mass Spectrom. 2017;31(22):1915–1926.

18. Arul AB, Robinson RAS. Sample Multiplexing Strategies in Quantitative Proteomics. Anal Chem. 2019;91(1):178–189. doi:10.1021/acs.analchem.8b05626

19. Cheung TK, Lee CY, Bayer FP, McCoy A, Kuster B, Rose CM. Defining the carrier proteome limit for single-cell proteomics. Nat Methods. Published online 2021. doi:10.1038/s41592-020-01002-5

20. Ye Z, Batth TS, Rüther P, Olsen J V. A deeper look at carrier proteome effects for single- cell proteomics. Commun Biol. 2022;5(1):150. doi:10.1038/s42003-022-03095-4

21. Biringer RG, Horner JA, Viner R, Hühmer AFR, Specht A. Quantitation of TMT-Labeled Peptides Using Higher-Energy Collisional Dissociation on the Velos Pro Ion Trap Mass Spectrometer. Thermo Fish Sci. Published online 2011.

22. Paulo JA, O’Connell JD, Gygi SP. A Triple Knockout (TKO) Proteomics Standard for Diagnosing Ion Interference in Isobaric Labeling Experiments. J Am Soc Mass Spectrom. Published online 2016. doi:10.1007/s13361-016-1434-9

23. Thompson A, Wölmer N, Koncarevic S, et al. TMTpro: Design, Synthesis, and Initial Evaluation of a Proline-Based Isobaric 16-Plex Tandem Mass Tag Reagent Set. Anal Chem. Published online 2019. doi:10.1021/acs.analchem.9b04474

24. Li J, Cai Z, Bomgarden RD, et al. TMTpro-18plex: The Expanded and Complete Set of TMTpro Reagents for Sample Multiplexing. J Proteome Res. 2021;20(5):2964–2972. doi:10.1021/acs.jproteome.1c00168

25. Michalski A, Damoc E, Lange O, et al. Ultra high resolution linear ion trap orbitrap mass spectrometer (orbitrap elite) facilitates top down LC MS/MS and versatile peptide fragmentation modes. Mol Cell Proteomics. Published online 2012. doi:10.1074/mcp.O111.013698

26. O’Connell JD, Paulo JA, O’Brien JJ, Gygi SP. Proteome-Wide Evaluation of Two Common Protein Quantification Methods. J Proteome Res. 2018;17(5):1934–1942. doi:10.1021/acs.jproteome.8b00016

27. Shen X, Shen S, Li J, et al. IonStar enables high-precision, low-missing-data proteomics quantification in large biological cohorts. Proc Natl Acad Sci. Published online 2018. doi:10.1073/pnas.1800541115

28. Gygi JP, Yu Q, Navarrete-Perea J, Rad R, Gygi SP, Paulo JA. Web-Based Search Tool for Visualizing Instrument Performance Using the Triple Knockout (TKO) Proteome Standard. J Proteome Res. 2019;18(2):687–693. doi:10.1021/acs.jproteome.8b00737

29. Hsiao CJ, Tung P, Blischak JD, et al. Characterizing and inferring quantitative cell cycle phase in single-cell RNA-seq data analysis. Genome Res. 2020;30(4):611–621. doi:10.1101/gr.247759.118

30. Barron M, Li J. Identifying and removing the cell-cycle effect from single-cell RNA- Sequencing data. Sci Rep. 2016;6(1):33892. doi:10.1038/srep33892

31. Brunner A-D, Thielert M, Vasilopoulou C, et al. Ultra-high sensitivity mass spectrometry quantifies single-cell proteome changes upon perturbation. Mol Syst Biol. 2022;18(3):e10798. doi:https://doi.org/10.15252/msb.202110798

32. Kelly V, al-Rawi A, Lewis D, Kustatscher G, Ly T. Low Cell Number Proteomic Analysis Using In-Cell Protease Digests Reveals a Robust Signature for Cell Cycle State Classification. Mol Cell Proteomics. 2022;21(1). doi:10.1016/j.mcpro.2021.100169

33. Orsburn BC. Evaluation of the Sensitivity of Proteomics Methods Using the Absolute Copy Number of Proteins in a Single Cell as a Metric. Proteomes. 2021;9(3). doi:10.3390/proteomes9030034

34. Taus T, Köcher T, Pichler P, et al. Universal and confident phosphorylation site localization using phosphoRS. J Proteome Res. Published online 2011. doi:10.1021/pr200611n

35. Kong AT, Leprevost F V., Avtonomov DM, Mellacheruvu D, Nesvizhskii AI. MSFragger: Ultrafast and comprehensive peptide identification in mass spectrometry-based proteomics. Nat Methods. Published online 2017. doi:10.1038/nmeth.4256

36. Dorfer V, Pichler P, Stranzl T, et al. MS Amanda, a universal identification algorithm optimized for high accuracy tandem mass spectra. J Proteome Res. Published online 2014. doi:10.1021/pr500202e

37. Kapp EA, Schütz F, Connolly LM, et al. An evaluation, comparison, and accurate benchmarking of several publicly available MS/MS search algorithms: Sensitivity and specificity analysis. Proteomics. Published online 2005. doi:10.1002/pmic.200500126

38. Käll L, Canterbury JD, Weston J, Noble WS, MacCoss MJ. Semi-supervised learning for peptide identification from shotgun proteomics datasets. Nat Methods. Published online 2007. doi:10.1038/nmeth1113

39. Liu SY, Ikegami K. Nuclear lamin phosphorylation: an emerging role in gene regulation and pathogenesis of laminopathies. Nucleus. 2020;11(1):299–314. doi:10.1080/19491034.2020.1832734

40. Kochin V, Shimi T, Torvaldson E, et al. Interphase phosphorylation of lamin A. J Cell Sci. 2014;127(Pt 12):2683–2696. doi:10.1242/jcs.141820

41. Garcia BA, Mollah S, Ueberheide BM, et al. Chemical derivatization of histones for facilitated analysis by mass spectrometry. Nat Protoc. 2007;2(4):933–938. doi:10.1038/nprot.2007.106

42. Jiang G, Li C, Lu M, Lu K, Li H. Protein lysine crotonylation: past, present, perspective. Cell Death Dis. 2021;12(7):703. doi:10.1038/s41419-021-03987-z

43. Wan J, Liu H, Chu J, Zhang H. Functions and mechanisms of lysine crotonylation. J Cell Mol Med. 2019;23(11):7163–7169. doi:10.1111/jcmm.14650

44. Wu Q, Li W, Wang C, et al. Ultradeep Lysine Crotonylome Reveals the Crotonylation Enhancement on Both Histones and Nonhistone Proteins by SAHA Treatment. J Proteome Res. 2017;16(10):3664–3671. doi:10.1021/acs.jproteome.7b00380

45. Singh C, Zampronio CG, Creese AJ, Cooper HJ. Higher energy collision dissociation (HCD) product ion-triggered electron transfer dissociation (ETD) mass spectrometry for the analysis of N-linked glycoproteins. J Proteome Res. Published online 2012. doi:10.1021/pr300257c

46. Stadlmann J, Taubenschmid J, Wenzel D, et al. Comparative glycoproteomics of stem cells identifies new players in ricin toxicity. Nature. Published online 2017. doi:10.1038/nature24015

47. Stadlmann J, Hoi DM, Taubenschmid J, Mechtler K, Penninger JM. Analysis of PNGase F-Resistant N-Glycopeptides Using SugarQb for Proteome Discoverer 2.1 Reveals Cryptic Substrate Specificities. Proteomics. Published online 2018. doi:10.1002/pmic.201700436

48. Federspiel JD, Greco TM, Lum KK, Cristea IM. Hdac4 Interactions in Huntington’s Disease Viewed Through the Prism of Multiomics *. Mol Cell Proteomics. 2019;18(8):S92–S113. doi:10.1074/mcp.RA118.001253

49. Xue JY, Zhao Y, Aronowitz J, et al. Rapid non-uniform adaptation to conformation-specific KRAS(G12C) inhibition. Nature. Published online 2020. doi:10.1038/s41586-019-1884-x

50. Drosten M, Barbacid M. Targeting the MAPK Pathway in KRAS-Driven Tumors. Cancer Cell. Published online 2020. doi:10.1016/j.ccell.2020.03.013

51. Santana-Codina N, Chandhoke AS, Yu Q, et al. Defining and Targeting Adaptations to Oncogenic KRASG12C Inhibition Using Quantitative Temporal Proteomics. Cell Rep. Published online 2020. doi:10.1016/j.celrep.2020.03.021

52. Leduc A, Huffman RG, Slavov N. Droplet sample preparation for single-cell proteomics applied to the cell cycle. bioRxiv. Published online January 1, 2021:2021.04.24.441211. doi:10.1101/2021.04.24.441211

53. Williams SM, Liyu A V, Tsai C-F, et al. Automated Coupling of Nanodroplet Sample Preparation with Liquid Chromatography–Mass Spectrometry for High-Throughput Single- Cell Proteomics. Anal Chem. 2020;92(15):10588–10596. doi:10.1021/acs.analchem.0c01551

54. Zhu Y, Clair G, Chrisler WB, et al. Proteomic Analysis of Single Mammalian Cells Enabled by Microfluidic Nanodroplet Sample Preparation and Ultrasensitive NanoLC-MS. Angew Chemie Int Ed. 2018;57(38):12370–12374. doi:https://doi.org/10.1002/anie.201802843

55. Schoof EM, Furtwängler B, Üresin N, et al. Quantitative single-cell proteomics as a tool to characterize cellular hierarchies. Nat Commun. 2021;12(1):3341. doi:10.1038/s41467-021-23667-y

56. Huffman RG, Chen A, Specht H, Slavov N. DO-MS: Data-Driven Optimization of Mass Spectrometry Methods. J Proteome Res. Published online 2019. doi:10.1021/acs.jproteome.9b00039

57. Boekweg H, Van Der Watt D, Truong T, et al. Features of Peptide Fragmentation Spectra in Single-Cell Proteomics. J Proteome Res. 2022;21(1):182–188. doi:10.1021/acs.jproteome.1c00670

58. Stopfer LE, Conage-Pough JE, White FM. Quantitative Consequences of Protein Carriers in Immunopeptidomics and Tyrosine Phosphorylation MS2 Analyses. Mol Cell Proteomics. 2021;20:100104. doi:https://doi.org/10.1016/j.mcpro.2021.100104

59. Lombard-Banek C, Choi SB, Nemes P. Chapter Thirteen - Single-cell proteomics in complex tissues using microprobe capillary electrophoresis mass spectrometry. In: Allbritton NL, Kovarik MLBT-M in E, eds. Enzyme Activity in Single Cells. Vol 628. Academic Press; 2019:263–292. doi:https://doi.org/10.1016/bs.mie.2019.07.001

60. Mund A, Coscia F, Kriston A, et al. Deep Visual Proteomics defines single-cell identity and heterogeneity. Nat Biotechnol. Published online 2022. doi:10.1038/s41587-022-01302-5

61. Hata AN, Shaw AT. Resistance looms for KRASG12C inhibitors. Nat Med. Published online 2020. doi:10.1038/s41591-020-0765-z

62. Ye X, Chan KC, Waters AM, et al. Comparative proteomics of a model MCF10A- KRasG12V cell line reveals a distinct molecular signature of the KRasG12V cell surface. Oncotarget. Published online 2016. doi:10.18632/oncotarget.13566

63. Scheltema RA, Hauschild J-P, Lange O, et al. The Q Exactive HF, a Benchtop Mass Spectrometer with a Pre-filter, High-performance Quadrupole and an Ultra-high-field Orbitrap Analyzer. Mol Cell Proteomics. Published online 2014. doi:10.1074/mcp.M114.043489

64. Jenkins C, Orsburn BC. Diagnostic Ion Data Analysis Reduction (DIDAR) allows rapid quality control analysis and filtering of multiplexed single cell proteomics data. bioRxiv. Published online January 1, 2022:2022.02.22.481489. doi:10.1101/2022.02.22.481489

